# The basolateral amygdala-anterior cingulate pathway contributes to depression and its comorbidity with chronic pain

**DOI:** 10.1101/2022.08.09.503276

**Authors:** Léa J Becker, Clémentine Fillinger, Robin Waegaert, Pierre Hener, Beyza Ayazgok, Muris Humo, Sarah H Journée, Meltem Karatas, Laetitia Degiorgis, Marie des Neiges Santin, Mary Mondino, Michel Barrot, El Chérif Ibrahim, Gustavo Turecki, Raoul Belzeaux, Pierre Veinante, Laura A Harsan, Sylvain Hugel, Pierre-Eric Lutz, Ipek Yalcin

## Abstract

While depression and chronic pain are frequently comorbid, underlying neuronal circuits, and their relevance for the understanding of psychopathology, remain poorly defined. Here we show in mice that hyperactivity of the neuronal pathway linking the basolateral amygdala to the anterior cingulate cortex is essential for chronic pain-induced depression. In naive animals, we demonstrate that activation of this pathway is sufficient to trigger depressive-like behaviors, as well as transcriptomic alterations that recapitulate core molecular features of depression in the human brain. These alterations notably impact gene modules related to myelination and the oligodendrocyte lineage. Among these, we show that Sema4a, a hub gene significantly upregulated in both mice and humans in the context of altered mood, is necessary for the emergence of depressive-like behaviors. Overall, these results place the BLA-ACC pathway at the core of pain and depression comorbidity, and unravel the role of impaired myelination and *Sema4a* in mood control.

## Main

Major depressive disorder (MDD) and chronic pain are long-lasting detrimental conditions that significantly contribute to the worldwide burden of disease^1, 2^. These two pathologies are highly comorbid, which results in increased disability and poorer prognosis compared to either condition alone^3, 4^. Despite their high prevalence and co-occurrence, available treatments remain ineffective, urging for a better understanding of the pathophysiology of this comorbidity.

The frontal cortex, particularly the anterior cingulate cortex (ACC), is at the center of emotional and pain processing^5^. A large body of evidence documents major alterations in ACC neuronal activity in patients with either chronic pain or MDD, as well as in rodent models of each condition^6–9^. Among other findings, our group previously showed that a lesion^10^ or the optogenetic inhibition^11^ of the ACC alleviates anxiodepressive-like consequences of neuropathic pain in mice, while activation of this structure is sufficient to trigger emotional dysfunction in naive animals. These results highlight a critical role of the ACC in the comorbidity between pain and MDD. However, the initial mechanisms priming ACC dysfunction remain poorly understood.

MDD originates from alterations affecting both the subcortical processing of external and internal stimuli, and their integration into perceived emotions by higher-level cortical structures^12^. It is, therefore, critical to understand how subcortical inputs to the ACC contribute to the emergence of emotional dysfunction. In extension to other afferences, the anterior part of the basolateral nucleus of the amygdala (BLA) shows dense, direct and reciprocal connections with the areas 24a/b of the ACC^13, 14^, which have been poorly studied in animal models of pain and mood disorders. The BLA plays a critical role in emotional processes, since its neurons encode stimuli with a positive valence, such as rewards^15, 16^, as well as those with a negative valence, including fear^16^ or pain states^17^. In humans, neuroimaging studies have consistently found that depressed patients^7, 18^ and individuals with chronic pain^6, 19^ exhibit pathological ACC and BLA hyperactivity.

In this context, we hypothesized that the neuronal pathway linking the BLA and the ACC might represent a core substrate underlying the comorbidity between pain and MDD. First, using retrograde tracing and neuronal activity markers, as well as rodent brain functional magnetic resonance imaging (MRI), we demonstrate that this pathway is hyperactive when chronic pain triggers depressive-like behaviors. Then, by optogenetically manipulating this circuit, we show that its hyperactivity is both necessary for chronic pain-induced depressive-like (CPID) behaviors, and sufficient to trigger similar deficits in naive animal, in the absence of chronic pain. We next characterize transcriptomic changes occurring in the ACC when BLA-ACC hyperactivity triggers mood dysfunction, and find that they strikingly resemble the molecular blueprint of depression in humans. These results, which notably include alterations affecting oligodendrocytes and myelination, are synergistic with our mouse imaging data showing altered tissue anisotropy along the BLA-ACC pathway, and establish the translational relevance of our BLA-ACC optogenetic model of MDD. Finally, we leverage gene network approaches to prioritize *Semaphorin 4A* (*Sema4A)* as a hub gene that is significantly upregulated in both mice and men in the context of altered mood. Using a gene knockdown approach, we demonstrate that, following optogenetic BLA-ACC stimulation, upregulation of *Sema4A* is necessary for the emergence of depressive-like behaviors. Overall, these results uncover the BLA-ACC pathway as a core substrate of pain and MDD comorbidity.

## Results

### Chronic neuropathic pain induces hyperactivity in the BLA-ACC pathway

To confirm the anatomical connection between the BLA and ACC, we first injected the retrograde tracer cholera toxin subunit B (CTB) into the areas 24a/b of the ACC (**Fig. 1a**). Consistent with previous reports, strongly labelled cell bodies were found in the BLA (**Fig. 1b**)^13, 20, 21^. Next, considering that we^11^, and others^5, 8, 22^, have found that the ACC is hyperactive when chronic pain triggers depressive-like behaviors, we wondered whether the BLA is similarly affected. To address this question, we quantified the immediate early gene c-Fos in our well-characterized CPID model^23, 24^. In this model, peripheral nerve injury leads to immediate and long-lasting mechanical hypersensitivity, with delayed anxiodepressive-like behaviors (significant at 7 weeks post-operation, PO; **Fig. 1c-f**). In our previous study, increased c-Fos immunoreactivity was found in the ACC at 8 weeks PO^25^. Here, a similar c-Fos increase was found in the BLA (**Extended Data Fig. 1a-b**). This indicates concurrent neuronal hyperactivity of the 2 structures when anxiodepressive-like behaviors are present. Next, we studied whether these c-Fos positive cells in the BLA directly innervate the ACC (**Fig. 1g**, right BLA, ipsilateral to nerve injury, **Extended Data Fig. 1c**, left BLA, contralateral to nerve injury). The retrograde tracer fluorogold (FG) was injected into the ACC of neuropathic animals, and we quantified its co-localization with c-Fos in the BLA at 8 weeks PO (**Fig. 1c**). Similar numbers of BLA neurons projecting to the ACC (FG+) were found in both sham and CPID groups, indicating that neuropathic pain does not modify the number of neurons in this pathway (**Fig. 1h**). An increase in the total number of c-Fos+ cells was observed in the right BLA of CPID mice, compared to sham (**Fig. 1i**), confirming results from our previous cohort (**Extended Data Fig. 1a-b**). Importantly, neuronal hyperactivity was, notably, observed in ACC-projecting neurons in the CPID group, as evidenced by an increase in cells positive for both FG and c-Fos (**Fig. 1j; Extended Data Fig. 1d**) in the BLA ipsilateral to the nerve injury (see **Extended Data Fig. 1e-g** for contralateral data). While a similar lateralization was already reported in the central amygdala during chronic pain^26, 27^, our results extend these findings to the BLA in the context of CPID. Finally, as a complementary strategy to assess the connectivity of the ACC and BLA, we took advantage of brain imaging data recently generated using resting-state functional Magnetic Resonance Imaging (rsfMRI). Consistent with above histological analyses, we observed that functional connectivity between the 2 structures was enhanced at 8, but not 2, weeks PO in CPID animals compared to sham controls (**Fig. 1k; Extended Data Fig. 1h-i** for behavioral characterization). Altogether, these results indicate that chronic pain induces hyperactivity of BLA neurons projecting to the ACC.

**Figure 1.**
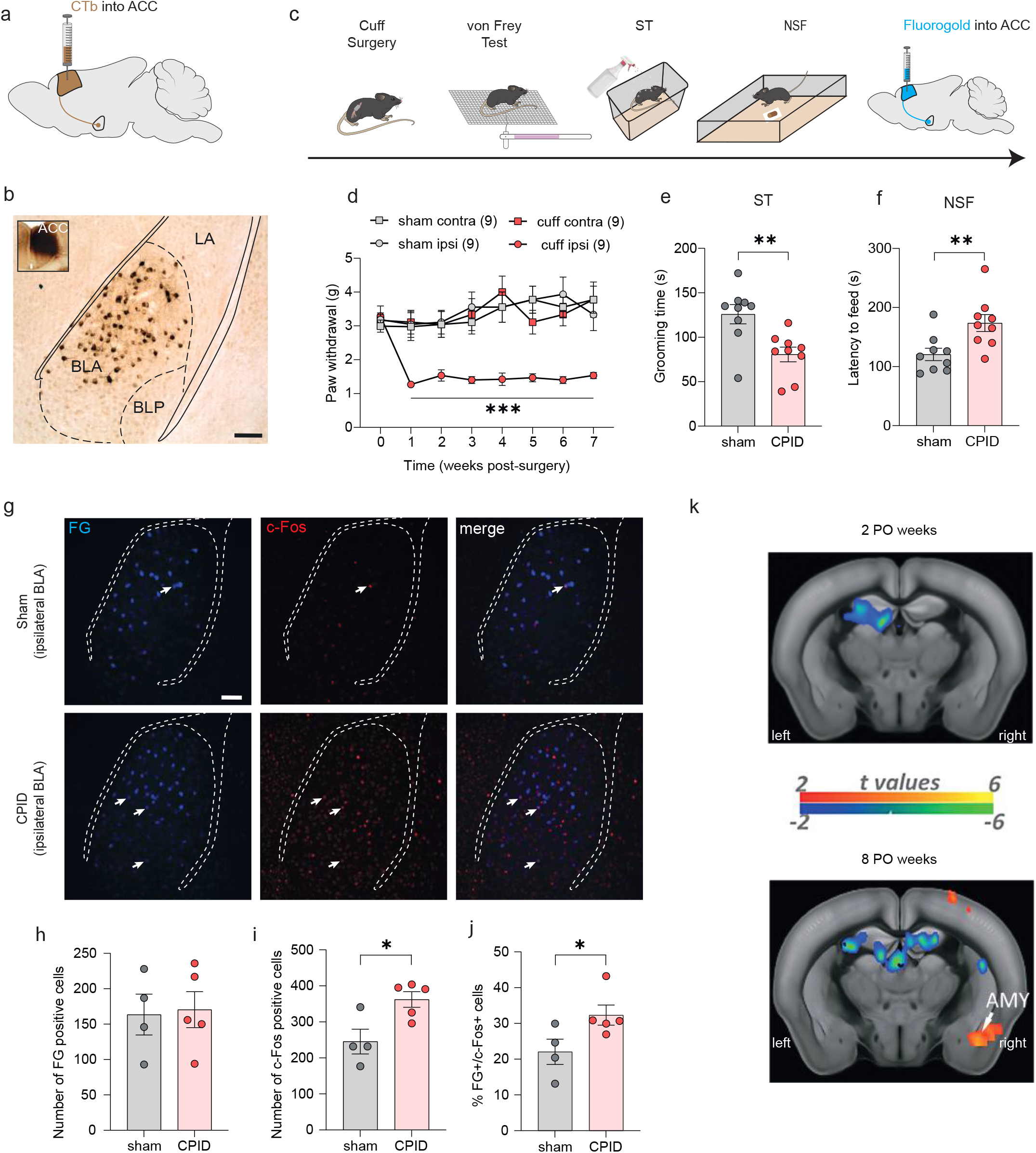
Chronic pain induced-depression (CPID) triggers hyperactivity in the BLA neurons projecting to the ACC and increases the functional connectivity between the ACC and the BLA. **a.** Illustration of the retrograde tracing strategy, with the injection of the cholera toxin B subunit (CTB) into the mouse ACC. **b.** Representative image of retrogradely labelled cell bodies in the BLA. Scale bar=100µm **c.** Experimental design for quantification of neuronal activity of BLA neurons projecting to the ACC in the CPID model. Peripheral nerve injury is induced by implanting a cuff around the main branch of sciatic nerve. Mechanical threshold is evaluated using von Frey filaments, and anxiodepressive-like behaviors using splash test (ST) and novelty suppressed feeding test (NSF). Fluorogold is injected into the ACC to label the afferent neurons. **d-f**. Peripheral nerve injury induced an ipsilateral long-lasting mechanical hypersensitivity (d; sham: n=9; cuff: n=9; F_(21,224)_=2.710; p<0.0001; post-hoc weeks 1-7 p<0.05), decreases grooming behavior in the splash test (e; sham: n=9; 125.90±10.80; CPID: n=9; 80.67±8.22; p=0.0042) and increases latency to feed in novelty suppressed feeding test (NSF) (f; sham: n=9; 120.7±10.49; CPID: n=9; 173.7±14.38; p=0.0089). **g.** Representative fluorescence images showing cells positive for fluorogold (FG+, left panel), c-Fos (c-Fos+, middle panel), or co-labelled (right panel) cells. **h-j**. Quantification of FG+, c-Fos+ cells and their co-localization revealed that, 8 weeks after peripheral nerve injury, the number of FG+ cells was not altered (panel h; sham: 163.5±28.85; CPID: 170.6±25.43; p=0.3651), while c-Fos+ (panel i; sham: 245.5±34.37; CPID: 362.4±21.62; p=0.0238) and FG+/c-Fos+ cells (panel j; sham 22.06%±3.55; CPID: 32.32%±2.81; p=0.0159) were increased in the right BLA (sham: n=4; CPID: n=5). Scale bar=100µm. **k.** Inter-groups statistical comparison results showed increased functional connectivity between the ACC and the amygdala (AMY) at 8 (bottom image) but not 2 post-operative (PO) weeks (upper image). FWER corrected at cluster level for p<0.05. Data are represented as mean±SEM. *p<0.05; **p<0.01. 2-Way ANOVA repeated measures (Time x Surgery; VF); unpaired t-test (ST and NSF); one-tailed Mann-Whitney test (FG, c-Fos quantification). CPID: Chronic pain induced depression; contra: contralateral; ipsi: ipsilateral, LA: lateral nucleus of the amygdala; AMY: amygdala. PO: post-operative.

### Optogenetic inhibition of the BLA-ACC pathway prevents CPID

We next hypothesized that this hyperactivity may be responsible for emotional dysfunction. To address this, we inhibited this pathway using an AAV5-CamKIIa-ArchT3.0-EYFP vector injected bilaterally in the BLA (**Fig. 2a-b**). To characterize the effects of green light illumination on BLA neurons, we performed *ex vivo* electrophysiological recordings 6 weeks after viral injection (**Fig. 2c**). Patch-clamp recordings at the level of the BLA showed that light stimulation resulted in neuronal inhibition, proportional to light intensity, as indicated by outward currents recorded in voltage-clamp mode (**Fig. 2d-e**).

**Figure 2.**
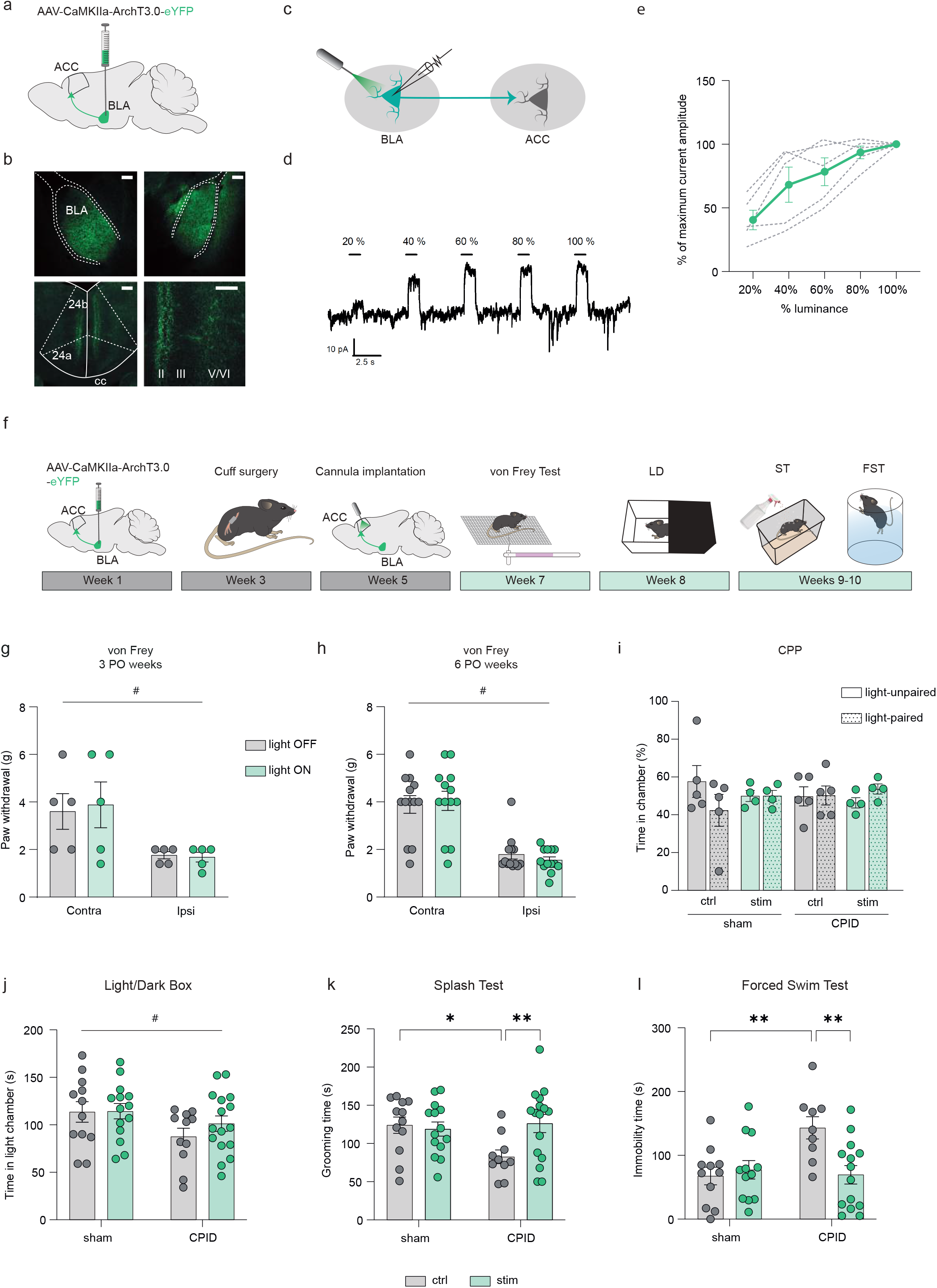
Optogenetic inhibition of the BLA-ACC pathway blocks CPID. **a**. Graphical representation of inhibitory AAV-CamkIIa-ArchT3.0-eYFP virus delivery to the mouse BLA for *ex-vivo* voltage-clamp recordings. **b**. Representative images of eYFP+ cell bodies in the BLA (upper panels) and eYFP+ axon terminals in the ACC (lower panels). Scale bars=100µm **c**. Graphical representation of the configuration for patch-clamp recording in the BLA **d**. Representative trace of the outward currents induced by optogenetic stimulation with increased luminance in a BLA neuron. **e**. Amplitude of currents induced by optogenetic stimulations of BLA neurons as a function of light stimulation intensity (green trace=mean; grey traces=individual responses). **f**. Graphical representation of the experimental design for *in-vivo* optogenetic inhibition of the BLA-ACC pathway, including bilateral virus delivery, peripheral nerve injury (cuff), cannula implantation, and behavioral testing. **g-h**. At 3 or 6 weeks after peripheral nerve injury, mechanical hypersensitivity was not affected by the inhibition of BLA-ACC pathway (ipsi vs contra; 3 PO weeks F_(1,4)_=7.752; p=0.0496; 6 PO weeks F_(1,12)_=55.80; p<0.0001; light-off vs light-on; 3 PO weeks F_(1,4)_=0.669; p=0.4592; 6 PO weeks F_(1,12)_=2.971; p=0.1104). **i.** Optogenetic inhibition of the BLA-ACC pathway did not induce a place preference at 6 weeks PO in sham and CPID animals for the chamber in which light was delivered (F_(3,13)_=0.153; p=0.9998). **j**. At 7 weeks PO, optogenetic inhibition of the BLA-ACC pathway applied 5 minutes before the light-dark test, did not affect the decreased time spent in the light chamber observed in nerve-injured animals (sham vs CPID: F_(1,49)_ = 4.703; p=0.035; ctrl vs stim: F_(1,49)=_0.634; p=0.43). **k**. At 8 weeks PO, optogenetic inhibition of BLA-ACC pathway during the splash test reversed the decreased grooming behavior observed in nerve-injured non-stimulated animals without having any effect on sham animals (F_(1,48)_=4.991; p=0.03; post-hoc: sham-ctrl (n=12) > CPID-ctrl (n=10); p<0.05; CPID-ctrl (n=10) < CPID-stim(n=16); p<0.05 sham-ctrl (n=12) = sham-stim (n=14)). **l**. At 8 weeks PO, optogenetic inhibition of BLA-ACC pathway applied 5 minutes before the FST, blocked the increased immobility time observed in nerve-injured non-stimulated animals without having any effect on sham animals (F_(1,42)_=7.539; p=0.008, post-hoc: sham-ctrl (n=11) > CPID-ctrl (n=9), p<0.05; CPID-ctrl (n=9) > CPID-stim (n=14), p<0.01; sham-ctrl (n=11) = sham-stim (n=12)). Data are represented as mean±SEM. #=main effect; *p<0.05; **p<0.01. Two-way ANOVA repeated measures (von Frey); Two-way ANOVA (Surgery x Stimulation; LD, ST and FST). PO: post-operative. 24a, 24b: areas 24a and 24b of the ACC; II, III, V/VI: cortical layers of the ACC; cc: corpus callosum.

To assess behavioral effects of inhibiting the BLA-ACC pathway in CPID, the same viral vector was injected bilaterally in the BLA, followed by implantation of an optic fiber in the ACC to specifically inhibit axon terminals coming from the BLA (**Fig. 2f**). Acute inhibition did not impact mechanical thresholds, measured using von Frey, in sham (**Extended Data Fig. 2a-b**) and nerve-injured animals at either 3 or 6 weeks PO (**Fig. 2g-h**). The BLA-ACC pathway is, therefore, not essential for mechanical hypersensitivity, consistent with what several groups reported when manipulating the whole ACC^10, 28–30^. Likewise, the BLA-ACC pathway does not drive ongoing pain, since we did not observe any significant conditioned place preference (CPP) during optogenetic inhibition (**Fig. 2i**). Inhibiting the whole ACC was however sufficient to induce CPP in our previous work^11^. Therefore, modulation of pain states by the ACC likely recruits other structures than the BLA^31^.

We next assessed the effect of inhibiting the BLA-ACC pathway on anxiodepressive-like behaviors. Optogenetic inhibition was applied just before (for light/dark, LD, and forced swim tests, FST) or during behavioral testing (splash test, ST). As expected, CPID animal displayed significantly higher anxiety-like behaviors in the LD at 7 weeks PO (**Fig. 2j**) and higher depressive-like behaviors at 8 weeks PO (**Fig. 2k-l**), in both the ST and FST^11, 23^. Inhibiting the BLA-ACC pathway had no impact in the LD test, suggesting that other pathways may control anxiety-like consequences of chronic pain. In contrast, BLA-ACC inhibition completely reversed pain-induced decreased grooming in the ST (**Fig. 2k**), and significantly decreased immobility in the FST (**Fig. 2l**), revealing potent antidepressant-like effects. No effect of optogenetic inhibition was observed in sham animals, indicating that these antidepressant-like effects selectively manifest in the context of chronic pain. As a more general measure of emotionality^32^, we also calculated z-scores for each animal across tests (ST, FST). Results indicated that CPID mice showed global emotional deficit compared to sham controls, as indicated by lower z-scores, an effect that was completely prevented by inhibition of the BLA-ACC pathway (**Extended Data Fig. S2c**). Overall, these results demonstrate that hyperactivity of BLA neurons targeting the ACC is necessary for the selective expression of chronic pain-induced depressive-like behaviors.

### Repeated activation of the BLA-ACC pathway triggers depressive-like behaviors in naive mice

We next determined whether BLA-ACC hyperactivity is sufficient to induce depressive-like behaviors in naive mice, in the absence of neuropathic pain. An AAV5-CamKIIa-ChR2-EYFP vector was injected bilaterally in the BLA, and patch-clamp recordings confirmed that blue light stimulation (wavelength: 475nm, pulse duration: 10ms; frequency: 10Hz) evoked inward currents in eYFP-expressing BLA neurons, as well as evoked excitatory postsynaptic currents in pyramidal ACC neurons (**Fig. 3a-h**). In the BLA, optogenetic stimulation of cell bodies induced inward currents, with a plateau reached at 40% of maximal light intensity, while pulsed 10Hz stimulation produced strong and stable currents (**Fig. 3b-d**). In the ACC, stimulating BLA axon terminals induced strong inward currents, with a plateau at 80% of maximal light (**Fig. 3g-h**), indicating activation of ACC neurons.

**Figure 3.**
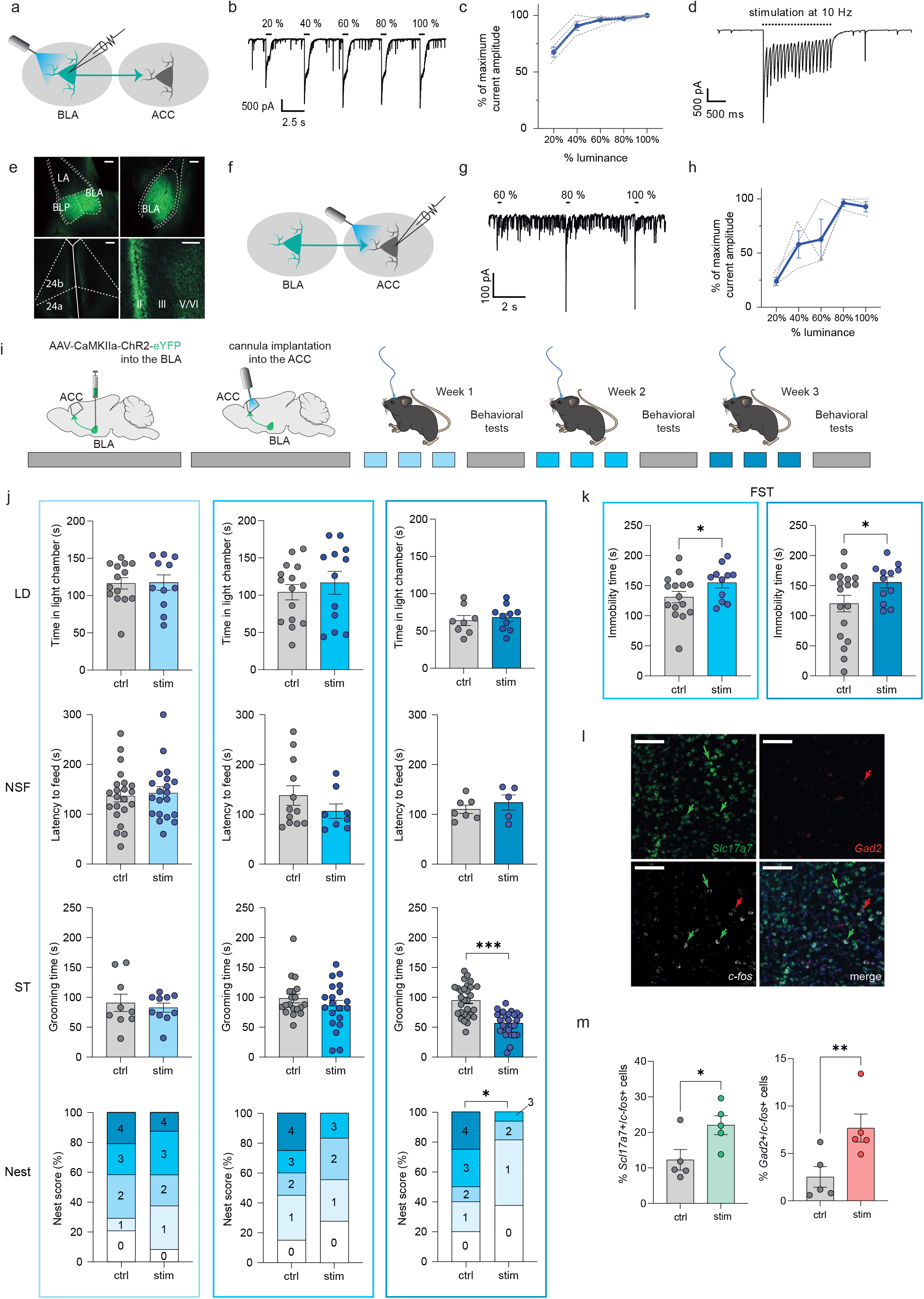
Repeated activation of the BLA-ACC pathway triggers depressive-like behaviors in naive mice. **a**. Graphical representation of the configuration for *ex-vivo* voltage-clamp recordings in the BLA. **b**. Representative trace of response of BLA neurons to 10Hz optogenetic activation showing that after an initial decrease in the amplitude of light-induced currents, a plateau is reached **c**. Representative trace of the outward currents induced by optogenetic stimulation with increased luminance in a BLA neuron. **d**. Amplitude of currents evoked by optogenetic stimulation of BLA neurons as a function of light stimulation intensity (blue trace=mean; grey traces=individual responses). **e**. Representative images of eYFP+ cell bodies in the BLA (upper panels) and eYFP+ axon terminals in the ACC (lower panels). Scale bars=100µm. **f**. Graphical representation of the configuration for *ex-vivo* voltage-clamp recordings in the ACC. **g**. Representative trace of the inward currents induced by optogenetic activation of BLA terminals within the ACC with increased luminance. **h**. Amplitude of currents evoked by optogenetic stimulations of BLA terminals recorded in ACC pyramidal neurons as a function of light stimulation intensities (blue trace=mean; grey traces=individual responses). **i**. Graphical representation of the experimental design for *in vivo* optogenetic activation of the BLA-ACC pathway, including bilateral virus delivery, cannula implantation and behavioral testing. **j**. Repeated activation of the BLA-ACC pathway did not induce anxiety-like behaviors in the LD (3 stim: ctrl: n=14; 116.4±7.57; stim: n=11; 117.4±10.31; p=0.94; 6 stim: ctrl: n=15; 104.0±10.31; stim: n=12; 116.7±15.37; p=0.49; 9 stim: ctrl: n=8; 64.0±6.73; stim: n=10; 68.0±5.34; p=0.64) and the NSF (3 stim: ctrl: n=22; 136.5±11.65; stim: n=20; 142.6±12.44; p=0.72; 6 stim: ctrl: n=12; 137.8±19.52; stim: n=8; 106.4±14.51; p=0.26; 9 stim: ctrl: n=7; 110.3±8.36; stim: n=5; 123.8±15.24; p=0.42) tests. While 3 and 6 sessions of optogenetic activation of the BLA-ACC pathway did not change the grooming behavior in the splash test (ST) (3 stim: ctrl: n=9; 90.78±14.42; stim: n=10; 82.70±7.52; p=0.62; 6 stim: ctrl: n=19; 98.16±7.37; stim: n=20; 86.10±8.7; p=0.30) and nest scores in nest test (3 stim: ctrl: n=24; stim: n=24; Chi-square=0.012; p=0.91; 6 stim: ctrl: n=20; stim: n=18; Chi-square=2.81; p=0.094), 9 stimulation decreased grooming time (ctrl : n=29; 94.79±5.02; stim: n=26; 56.81±3.96; p<0.0001) and nest quality (ctrl: n=20; stim: n=16; Chi-square=7.35; p=0.0067) in stimulated animals. **k**. The immobility time was increased after 6 and 9 stimulations of the BLA-ACC pathway in the forced swim test (FST; 6 stim: ctrl: n=15; 131.1±9.47; stim: n=11; 155.1±8.71; p=0.042; 9 stim: ctrl: n=18; 120.3±13.69; stim: n=12; 155.5±9.15; p=0.033). **l**. Representative images of *Slc17a7* (upper-left panel), *Gad2* (upper-right panel), *c-fos* (lower-left panel) mRNA expression and their co-localization (lower-right panel) in the ACC following BLA-ACC activation, measured by RNA scope. Scale bars=100µm. **m**. In the ACC, the proportion of *Slc17a7*+/*c-fos*+ cells (green) and the proportion of *Gad2*+/*c-fos*+ cells (red) was increased in stimulated animals (ctrl: n=5; stimulated : n=5; *Gad2*+/*c-fos*+: ctrl: 2.52±1.09; stim: 7.68±1.48, p=0.008; *Slc17a7*+/*c-fos*+: ctrl: 12.28±2.89; stim: 22.04±2.64, p=0.028). Data are represented as mean ± SEM. *p<0.05; **p<0.01; ***p<0.001. unpaired t-test (LD, NSF, ST); chi-square test for trend (Nest test); one-tailed Mann-Whitney test (*c-fos*/*Slc17a7*, *c-fos*/*Gad2* mRNA quantification). 24a, 24b: areas 24a and 24b of the ACC; II, III, V/VI: cortical layers of the ACC.

Having established this stimulation protocol, we explored its behavioral impact in naive mice. An optic fiber was implanted in the ACC, and blue light pulse stimulation applied for 20min (with parameters validated *ex vivo*). We first showed that a single session of stimulation was not sufficient to trigger any alterations in spontaneous locomotor activity, real-time place avoidance or anxiodepressive-like assays (**Extended Data Fig. 3a-d**). Thus, we next tested the effects of stimulations repeated over 3 consecutive days each week, during 3 weeks (**Fig. 3i-k**). Behavioral tests were performed following each block of 3 activating sessions (i.e. at the end of week 1, 2 or 3). No effects were found on locomotor activity (**Extended Data Fig. 3e**) or anxiety-like behaviors (**Fig. 3j**, LD/NSF), consistent with above CPID results indicating that optogenetic inhibition of the pathway did not rescue pain-induced anxiety-like behaviors. In contrast, depressive-like behaviors progressively emerged: (i) After the first 3 stimulations, no change was observed (**Fig. 3j**, ST/Nest test); (ii) After 6 stimulations, immobility in the FST significantly increased (**Fig. 3k**, left panel), with a tendency for decreased nest building, without significant effects in the ST; (iii) After 9 stimulations, emotional deficits further strengthened, as increased immobility in the FST (**Fig. 3k**, right panel) was accompanied by a significantly poorer nest score, along with decreased grooming in the ST (**Fig. 3j**). Of note, none of these effects remained detectable one week after the last stimulation (**Extended Data Fig. 3f-g**), indicating a reversible phenotype.

To determine which neuronal cell-types of the ACC are activated in this paradigm, we next quantified c-Fos expression (immunohistochemistry) and its co-localization (RNAscope) with markers of excitatory (glutamatergic, *Slc17a7*) and inhibitory (GABAergic, *Gad2*) neurons. Repeated optogenetic activation induced a strong increase in c-Fos expression (**Extended Data Fig. 3h**), predominantly in glutamatergic but also GABAergic cells (**Fig. 3l-m**, **Extended Data Fig. 3i-j**) of the ACC (24a/24b), while global numbers of *Gad2*+ or *Slc17a7*+ cells remained unaltered (**Extended Data Fig. 3k-m**). Altogether, these results show that repeated activation of the BLA-ACC pathway is sufficient to activate major neuronal cell types in the ACC, and triggers the progressive emergence of a depressive-like phenotype.

### Repeated activation of the BLA-ACC pathway produces transcriptional alterations similar to those observed in human depression

The need for repeated activations to produce behavioral effects suggests transcriptomic alterations within the ACC. To understand underlying molecular mechanisms, we used RNA-Sequencing to identify gene expression changes occurring in the ACC (**Extended Data Table S1**) after 9 stimulations (**Fig. 4a**), when behavioral deficits are maximal. We generated 2 animal cohorts (n=12 controls and n=10 stimulated mice in total) and, before harvesting ACC tissue, confirmed the development of depressive-like behaviors (**Fig. 4b**). Analysis of gene expression changes was conducted as described previously^33^ (see material and methods). Using Principal Component Analysis, robust differences were observed across groups at genome-wide level (**Extended Data Fig. 4a**). At nominal p-value (p<0.05), 2611 genes were significantly dysregulated in stimulated mice compared to controls, with 54 genes remaining significant after Benjamini-Hochberg correction for multiple testing (**Fig. 4c**). Over-Representation Analysis^34^ (ORA) uncovered alterations in Gene Ontology (GO) terms related to myelination, dendritic transport, neurogenesis and cytoskeleton (**Table S1; Extended Data Fig. 4b**).

**Figure 4.**
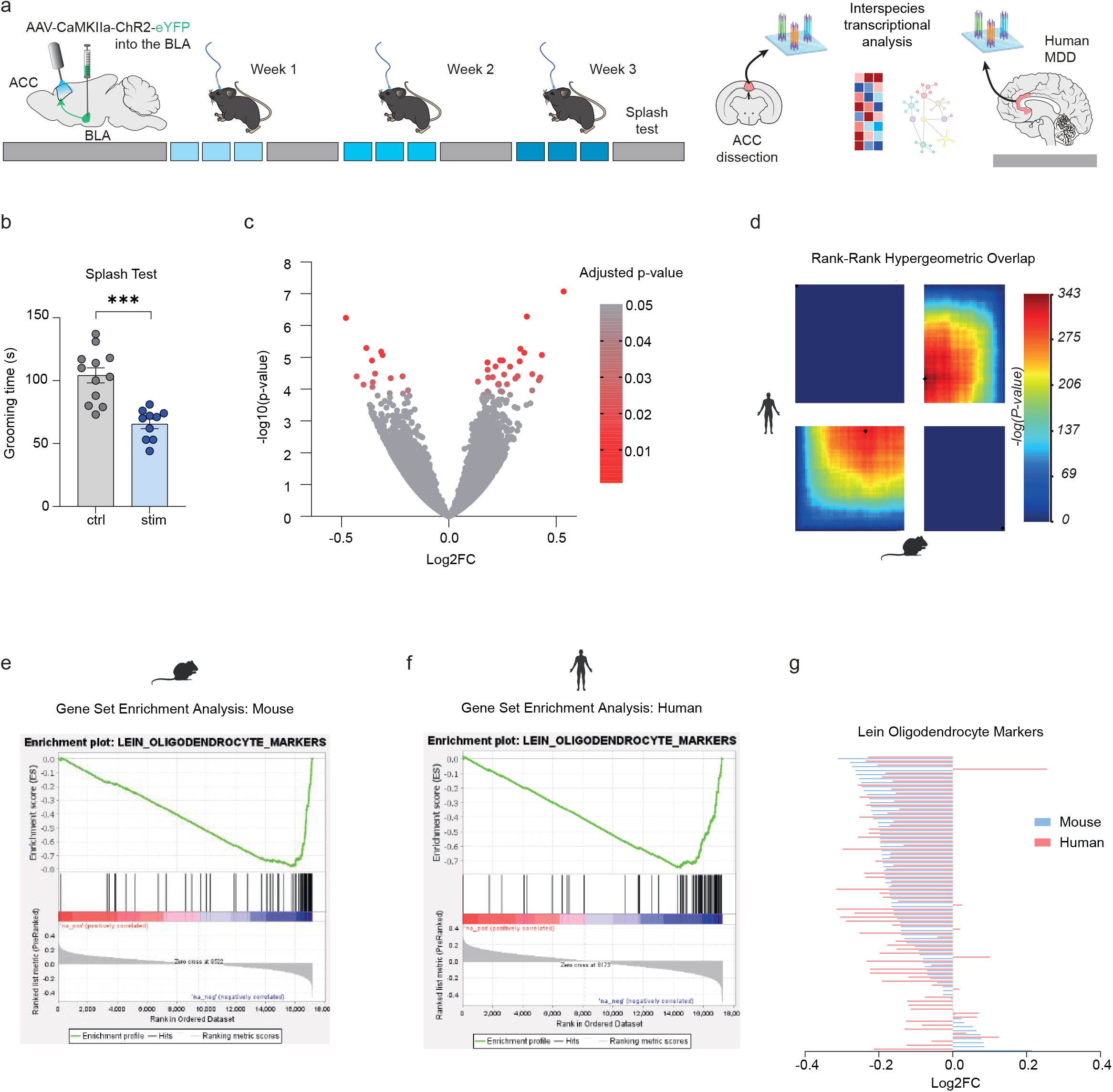
Repeated activation of the BLA-ACC pathway induces transcriptional alterations similar to those observed in human depressed patients. **a**. Graphical representation of experimental design, including virus delivery into the BLA, cannula implantation into the ACC, 9 sessions of optogenetic activation, ACC extraction in mice, and transcriptomic analysis in mice and human. **b**. 9 sessions of optogenetic activation of the BLA-ACC pathway decreases grooming behaviors in stimulated animals used for RNA-Sequencing (ctrl: n=12; 104.2±6.0; stim: n=10; 65.60±3.80; p<0.0001). **c**. Volcano plot showing the bidirectional distribution of the 2611 (6.9% of all genes) genes differentially expressed between stimulated (n=10) and control animals (n=12): red circles depict the 54 genes that showed a significant dysregulation after correction for multiple testing (padj<0.05). **d**. Rank-Rank Hypergeometric Overlap (RRHO2) unravels shared transcriptomic changes in the ACC across mice and men as a function of optogenetic stimulation (mouse) or a diagnosis of major depressive disorder (MDD). Levels of significance for the rank overlap between men and mice are color-coded, with a maximal Fisher’s Exact Test (FET) p=1.26E-153 for up-regulated genes (lower-left panel), and a maximal FET p=1.64E-127 for down-regulated genes (upper-right panel). **e-g** Gene set enrichment analysis (GSEA) revealed an enrichment for genes specifically expressed by oligodendrocytes and showing evidence of down-regulation as a function of optogenetic stimulation (e) or MDD diagnosis (f). The direction of the change correlated across mice and human (g; r²=0.15, p=0.0005). Behavioral data is represented as mean ± SEM, ***p<0.001, unpaired t-test.

To test relevance of these data to human depression, we compared them to our recent post-mortem study^35^. In the later, similar RNA-Seq was used to compare ACC tissue from individuals who died during a major depressive episode (n=26; **Extended Data Table S2**), and healthy individuals without any psychiatric history (n=24). Because only male mice were used in our paradigm, we reprocessed human data to restrict differential expression analysis to men (for a similar analysis using data generated from both sexes, see **Extended Data Fig. 5**). We then used 3 strategies to compare men and mice, focusing on 13 572 orthologues (see Methods). First, 398 genes were identified as differentially expressed (DEG, nominal p-value<0.05) in both species (common DEGs; **Extended Data Fig. 4c**). These represented 29.6% of all DEGs in mice (34.9% when considering both men and women; **Extended Data Fig. 5a-b**), indicating robust overlap between species (**Extended Data Fig. 4d**). ORA performed on common DEGs revealed enrichments in GO term related to neurogenesis, cytoskeleton or myelin sheath (**Extended Data Fig. 4e**). Second, for a more systematic threshold-free comparison, we used RRHO2, for Rank-Rank Hypergeometric Overlap^36^. This analysis uncovered large patterns of transcriptional dysregulation in similar directions as a function of MDD in men, and of optogenetic activation in mice. Indeed, strong overlaps were observed between species for both up-regulated and down-regulated genes (**Fig. 4d**), notably affecting pathways related to oligodendrocyte cell fate and myelination **(Table S3-4)**. Third, because results repeatedly pointed towards myelin, we used Gene Set Enrichment Analysis (GSEA, **Fig. 4e-g**) to interrogate a well-characterized list of 76 genes primarily expressed by oligodendrocytes, encompassing their major biological functions^37^. Among these, 48 (63%; **Fig. 4e**) were down-regulated in stimulated mice, and 57 in men with MDD (75%; **Fig. 4f**). This pattern of myelin down-regulation strongly correlated across species (**Fig. 4g**). Altogether, these 3 approaches (common DEG, RRHO2 and GSEA) converged to reveal significant dysregulation of myelination (see **Extended Data Fig. 4e-g**, **5c-e**).

### Gene ontology pathways affected in mice and men point towards altered oligodendrocyte function

To better capture the modular disorganization of gene expression in MDD, we then used weighted gene co-expression network analysis, WGCNA^38^, similar to our recent work^39^. Gene networks, constructed independently in each species, were composed of 25 and 29 gene modules in mice and humans, respectively. Using Fischer’s exact t-test, we found that the gene composition of 14 mice modules (56%) was significantly enriched in human modules, indicating their strong conservation (Benjamini-Hochberg, p<0.05; **Fig. 5a**; **Table S5**; see **Extended Data Fig. 5f; TableS6** when including women). To identify which modules are most significantly impacted, we computed correlations between optogenetic activation or MDD, and each module’s eigengene (a measure that summarizes co-expression). In humans, 12 module eigengenes significantly correlated with MDD; in mice, 6 modules associated with optogenetic stimulations (**Fig. 5b**, **Extended Data Fig. 5g**). Importantly, among these, 8/12 and 5/6 modules were also conserved across species. Therefore, human MDD and repeated optogenetic activation of the BLA-ACC pathway impact conserved gene modules in the ACC.

**Figure 5.**
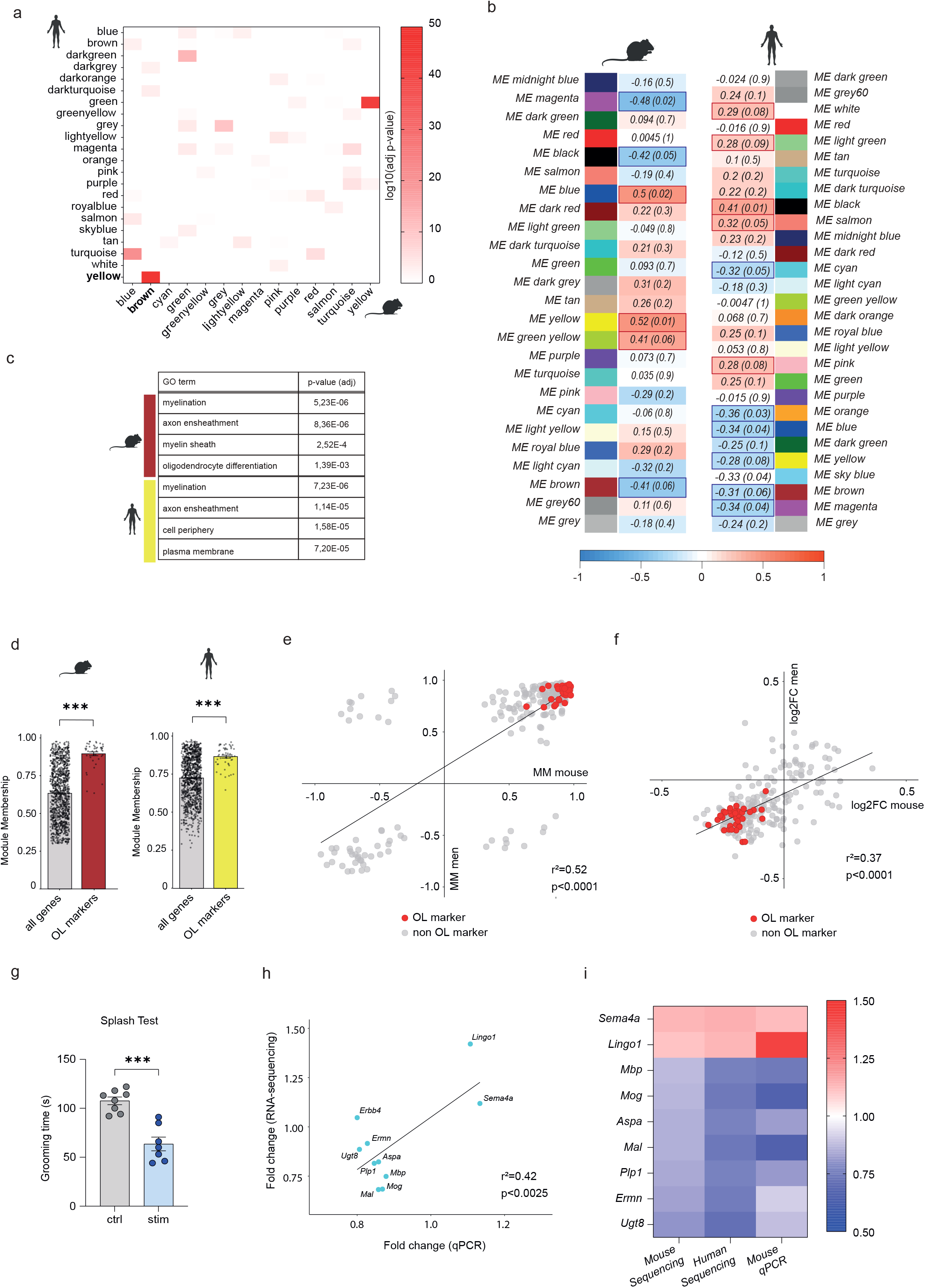
Gene-network analysis points toward an alteration of myelination and oligodendrocyte in mice and men. **a**. Heatmap representing the level of significance of overlaps between mice and men gene modules (measured using the FET). The highest overlap (p=8.36E-49) was obtained for the human/yellow and mouse/brown modules. **b**. WGCNA was used to analyze network and modular gene co-expression in the mouse and men ACC. The tables depict associations between individual gene modules and: i) optogenetic stimulation in mice (left panel), ii) or MDD diagnosis in men (right panel). Each row corresponds to correlations and p-values obtained against each module’s eigengene. **c**. Results of Gene Ontology enrichment analysis for human/yellow and mouse/brown module, with most significant enrichments corresponding to myelin related genes in both species. **d**. The absolute value of the module membership (MM) of oligodendrocytes (OL) markers is significantly higher than the MM of all the genes in each module for both mouse/brown (left panel; p-value<2.2E-16) or human/yellow (right panel; p-value<2.2E-16) modules. **e**. Linear regression of mouse (x-axis) and men (y-axis) MM computed by WGCNA, showing a significant positive correlation between Brown and Yellow modules gene ranking (r²=0.52, p=0.0001). Red dots indicate myelin and oligodendrocyte related genes/genes also present in the Lein-Oligodendrocytes-Markers data base. **f**. Linear regression of fold changes measured by RNA sequencing for men (x-axis) and mouse (y-axis), showing a significant positive correlation in the direction of the change in expression of the genes in Mouse/Brown and Men/Yellow modules (r²=0.37, p<0.0001). Red dots indicate myelin and oligodendrocyte related genes/genes also present in the Lein-Oligodendrocytes-Markers data base. **g**. Repeated activation of the BLA-ACC decreases grooming time in stimulated animals in splash test (ctrl n=8; 107.6±3.65; stim n=7; 63.57±6.96; p<0.0001). **h**. Linear regression of fold changes measured by RNA sequencing (x-axis) and qPCR (y-axis), showing a significant positive correlation between the two methods (r²=0.42, p=0.0025). **i**. Down-regulation of the myelin-related genes *(Mbp*, p=0.044; *Mog*, p=0.026; *Aspa*, p=0.078; *Mal*, p=0.038; *Plp1*, p 0.080; *Ermn*, p=0.180; *Ugt8*, p=0.133) and up-regulation of inhibitors of the myelination process (*Lingo1*, p=0.114; *Sema4a*, p=0.154) was consistently found across mice and men by RNA-Sequencing, and was validated by qPCR in mouse after 9 optogenetic stimulations. Behavioral data is represented as mean±SEM, ***p<0.0001, unpaired t-test.

We then conducted ORA of disorganized and conserved gene modules. Enriched GO terms were related to synaptic activity, mitochondria or RNA processing (**Extended Data Table S7**), consistent with previous studies of MDD^40–42^. Importantly, the 2 most strongly conserved modules (Men/Yellow and Mouse/Brown) again implicated myelin, with enrichments related to oligodendrocytes and myelination (**Fig. 5c**). To assess where myelin genes are located within these modules, we analyzed their module membership (MM), a measure of module centrality. Myelin-related genes displayed higher absolute values (i.e., higher centrality) compared to mean MM values among their host modules, in both species (**Fig. 5d**). Among the 235 genes belonging to the intersection between the 2 Men/Yellow and Mouse/Brown modules, 36 were myelin related **(Fig. 5e**, red dots), and a strong correlation between mouse and human MM was found, indicating that the same set of genes is centrally located among the 2 modules. Finally, we also observed a strong correlation in the directionality of gene expression changes across species, with a majority of down-regulated genes (**Fig. 5f**). Altogether, myelination deficiency, a critical feature of MDD pathophysiology in the ACC, was indeed modelled by our optogenetic paradigm.

We next validated RNA-sequencing results using microfluidic qPCR and a new mouse cohort generated with the same optogenetic protocol (n=8 control and 7 stimulated mice). Behavioral effects of stimulations were first confirmed (**Fig. 5g**), followed by dissection of the ACC tissue and analysis of the expression of most abundant myelin sheath proteins (*Plp1*, *Mal*, *Mog*, *Mag*, *Mbp*), enzymes involved in the synthesis of myelin lipids (*Aspa*, *Ugt8*), as well as positive (*Ermn*) and negative (*Sema4a*, *Lingo1*) regulators of myelination (**Fig. 5h-i**). Results strongly correlated with RNA sequencing data, with similar down-regulation of myelin sheath proteins, or enzymes for the synthesis of myelin lipids, as well as up-regulation of 2 well-known negative regulators of myelination, *Sema4a* and *Lingo1*.

Finally, as complementary approaches to document how these profound myelin gene expression changes translate at cellular and network levels, we used immunohistochemistry and brain imaging. In a first cohort, we assessed, in the ACC, the number of cells expressing Olig2, a transcription factor essential for proliferation and differentiation in the oligodendrocyte lineage. After 9 sessions of optogenetic stimulation, when depressive-like consequences are maximal (**Fig. 6a**), the number of Olig2-positive cells was significantly decreased (**Fig. 6b-c**), likely contributing to the decreased expression of myelin genes observed in bulk tissue. In another cohort (for behavioral validation, see **Extended Data Fig. 6a**), we performed MRI with DTI acquisition sequences, to analyze microstructural changes induced by optogenetic activation. Interestingly, stimulated animals displayed lower fractional anisotropy (FA) compared to controls in the ACC, amygdala and along the pathway connecting the two regions (**Fig. 6d**). This effect significantly correlated with increased depressive-like behavior (**Extended Data Fig. 6b** for behavioral validation). Since myelination is an important determinant of the structural connectivity assessed by DTI^43, 44^, these results reinforce the notion that repeated activation of the BLA-ACC pathway disrupts the transcriptional program and cell proliferation within the ACC oligodendrocyte lineage, leading to altered connectivity between the 2 structures.

**Figure 6.**
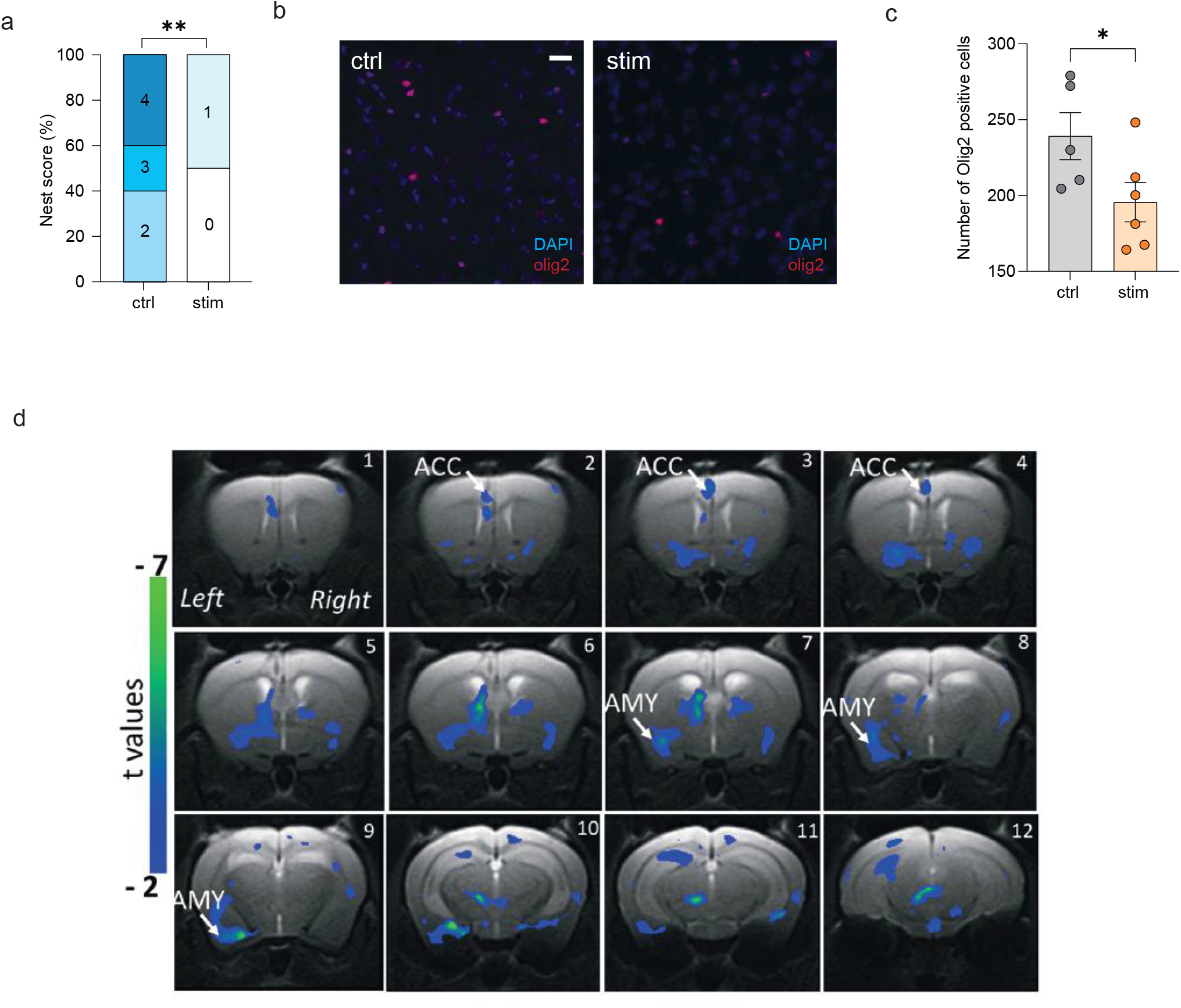
Repeated activation of the BLA-ACC pathway impairs myelination. **a**. Repeated activation of the BLA-ACC pathway decreased nest quality in stimulated animals (ctrl n=5; stim n=6; Chi-square=8.317; p=0.0039). **b.** Quantification of olig2 positive cells showed that repeated (9) stimulation of the BLA-ACC pathway decreased the number of olig2+ cells in the ACC (ctrl: n=5; 239.2±15.50; stim: n=6; 195.6±12.95; p=0.041). **c**. Representative fluorescence images showing cells that are olig2 positive (red) in non-stimulated (left panel) and stimulated mice (right panel). **d.** Representative coronal MRI images showing in blue the areas with significant decrease of FA along the left BLA-ACC pathway in stimulated animals compared to control group (ctrl: n=7; stim: n=6; GLM p<0.001 uncorrected) Data are represented as mean±SEM. *p<0.05; **p<0.01. chi-square test for trend (Nest test); one-tailed Mann-Whitney test (olig2 quantification).

### Sema4a is necessary for the emergence of optogenetically-induced depressive-like states

While the relationship between myelination and the expression of depressive-like behaviors has already been documented^45–49^, little is known about underlying molecular substrates. A few studies have investigated the effect of depleting myelin sheath protein^50^ or positive regulator of myelination^51^ on depressive-like behaviors in rodents. In contrast, upstream factors that prime myelin deficits and lead to behavioral dysregulation are unknown. Here, we focused on *Sema4a* because it is a potent inhibitor of myelination^52, 53^ that was recently associated with pathologies implicating white matter deficits^54, 55^, and was strongly up-regulated in our optogenetic paradigm (**Fig. 5i**).

To enable blocking the over-expression of *Sema4a* in our model, we first established a knock-down (KD) approach. Three different sh-RNAs (#108, 576, 791; **Fig. 7a**) targeting exon 15 of *Sema4a* were designed, packed into AAV plasmids with mCherry reporter, and transfected in HEK cells over-expressing the targeted *Sema4a* exon, in fusion with eGFP. Among these, we prioritized shRNA-791 as it yielded the most profound KD, as shown by a near complete loss of eGFP signal at 2 days post-transfection (**Fig. 7b**). To characterize *in vivo* efficiency of shRNA-791, it was then packaged into an AAV vector and injected in the ACC of adult mice, followed by qPCR quantification of *Sema4a* 6 weeks later (**Fig. 7c-d**). Compared with a control vector expressing the mCherry and a scrambled shRNA, the AAV-shRNA-791 achieved a 62% reduction of *Sema4a* expression, demonstrating robust efficacy.

**Figure 7.**
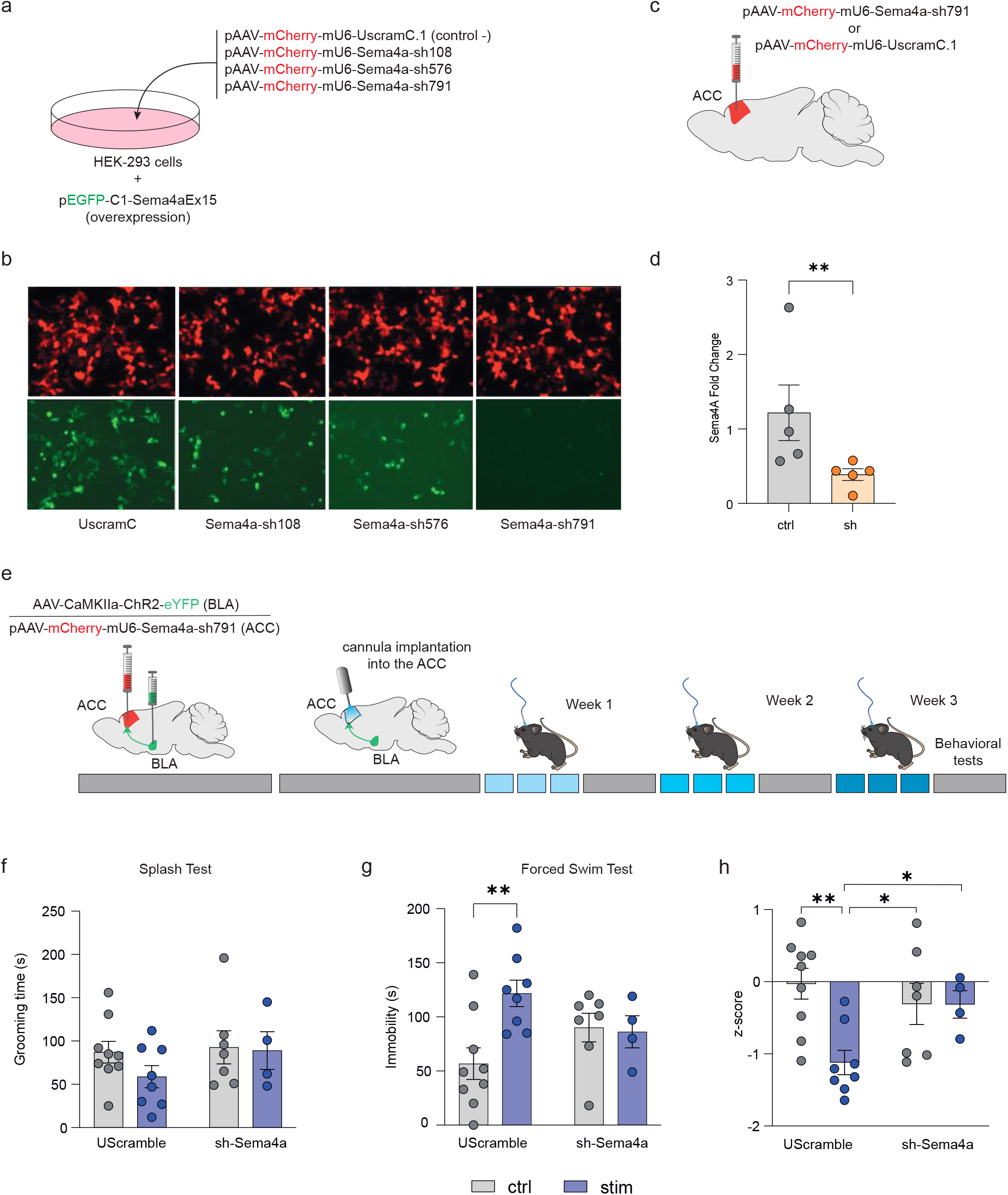
Semaphorin-4A is essential for the depressive-like behaviors induced by the activation of the BLA-ACC pathway. **a**. Graphical representation of the experimental design for AAV validation *in-vitro*. **b**. Representative fluorescence images showing HEK-293 cells expressing the mouse gene for *Sema4A* (green) and the transfected AAV (red) demonstrating the efficiency of the pAAV-mCherry-mU6-Sema4a-sh791 to knock-down *Sema4a*. **c**. Graphical representation of the bilateral virus injection in the ACC for *in-vivo* validation of the selected sh-RNA. **d.** qPCR analysis showing a down-regulation of *Sema4a* expression level in the ACC of mice injected with Sema4a-sh791 (ctrl: n=5; 1.22±0.37; stim: n=5; 0.39±0.07; p=0.0079). **e**. Graphical representation of the experimental design, including bilateral virus delivery in the BLA (AAV5-CaMKIIa-ChR2(H134R)-EYFP) and the ACC (rAAV-mCherry-scrambleUsh or rAAV-mCherry-Sema4a-sh791), cannula implantation, optogenetic stimulation and behavioral testing. **f.** Grooming time in the ST was not affected by repeated activation of the BLA-ACC pathway or by knocking down *Sema4a* in the ACC (F_(1,24)_=0.56; p=0.46). **g**. Knocking down the ACC *Sema4a* prevents the development of depressive-like behaviors observed in the FST in animals with repeated activation of the BLA-ACC pathway (F_(1,24)_=5.68; p=0.029; post-hoc: Uscramble-Ctrl<Uscramble-Stim; p=0.0015; sh-Sema4a-Ctrl=sh-Sema4a-Stim; p=0.87). **h**. Knocking down the ACC *Sema4a* normalized the emotionality z-score (F_(1,24)_=5.12; p=0.033; post-hoc: Uscramble-Ctrl>Uscramble-Stim; p=0.0011; UScramble-Stim<sh-Sema4a-Ctrl; p=0.016; UScramble-Stim<sh-Sema4a-Stim; p=0.04; sh-Sema4a-Ctrl=sh-Sema4a-Stim; p=0.99). Data are represented as mean±SEM. *p<0.05; **p<0.01; one-tailed Mann-Whitney test (*Sema4A* quantification), Two-way ANOVA (Stimulation x Knock-Down; ST, FST and z-score).

Finally, we hypothesized that knocking-down *Sema4a* prior to optogenetic stimulations may prevent the emergence of depressive-like behaviors. A new cohort of mice (**Fig. 7e**) went through bilateral injections of the ChR2-expressing virus in the BLA, followed by bilateral injections of the AAV-shRNA-791 vector (or the Scrambled control) in the ACC, and, 2 weeks later, optogenetic cannula implantation in the ACC. Behavioral testing was performed after 9 stimulations over 3 weeks, corresponding to the 6-week time-point at which we documented shRNA-791 *in vivo* efficiency (**Fig. 7d**). In this new cohort, no significant effect was observed in the ST (**Fig. 7f**). In the FST (**Fig. 7g**), we detected a significant interaction between optogenetic stimulations and *Sema4a* KD, with a potent increase in depressive-like behaviors of stimulated mice that did not occur when *Sema4a* was knocked-down. Analysis of emotional reactivity across ST and FST tests further strengthened these results, as global emotional dysfunction induced by BLA-ACC activation was reversed by *Sema4a* KD (**Fig. 7h**). Of note, *Sema4a* KD had no effect in unstimulated mice across any tests, indicating that it is not sufficient to trigger emotional dysfunction in naive animals. Altogether, these results indicate that silencing *Sema4a* in the ACC prevents the emergence of emotional deficits when the BLA-ACC is repeatedly activated, demonstrating the critical role of this inhibitor of myelin function in mood control.

## Discussion

Given the complexity of emotional consequences of chronic pain, disentangling the circuitries involved in its different components is crucial for uncovering new therapeutic leads and strategies. The ACC is considered to play a pivotal role in these processes^10, 30, 56^. However, while its connectome has been robustly established using neuroanatomical^13, 14^ and imaging approaches, how it integrates in polysynaptic neuronal circuits that may differentially regulate mood and nociception is poorly understood^57^. To address this gap, here we focused on the BLA-ACC pathway, based on their strong reciprocal anatomical connections and well-known functions. By leveraging 2 optogenetic strategies for neuronal activation or inhibition in the mouse, we manipulated ACC inputs coming from the BLA. Our translational results uncover a critical role of this discrete pathway in mood, both in the context of chronic pain or in the absence of any neuropathy.

First, we found that inhibiting the BLA-ACC pathway blocks the expression of depressive-like consequences of chronic pain, without affecting anxiety-like behaviors, aversiveness, or mechanical hypersensitivity. Because acute inhibition is sufficient to produce this effect, depressive-like features seem to be mediated by ongoing hyperactivity of the pathway. In contrast, our previous work had shown that inhibition of CamKIIa+ cells in the ACC, regardless of their connectivity features, attenuated both anxiety and depressive-like consequences of chronic pain^11^. This suggests that distinct ACC inputs differentially contribute to various aspects of the pain experience, which is coherent with data from other groups^31, 58^. Accordingly, Gao et al showed that, in the sciatic nerve chronic constriction injury model, inhibiting projections from the ACC to the mesolimbic pathway (nucleus accumbens and ventral tegmental area) induces CPP, without affecting evoked pain^31^. In parallel, Hirschberg et al found that ACC inputs coming from the locus coeruleus (LC) are involved in anxiety-like and aversive consequences of pain^58^. Combined with ours, these results suggest functional segregation, with a LC-ACC-mesolimbic circuit preferentially involved in pain-induced aversion, while BLA and LC inputs targeting the ACC may predominantly mediate depressive- and anxiety-like consequences of chronic pain, respectively.

Second, we show that, in the absence of neuropathy, repeated but not acute BLA-ACC activation is sufficient to trigger depressive-like effects. The lack of detectable impact of this optogenetic manipulation on anxiety-like responses further strengthens the aforementioned notion that ACC inputs coming from the BLA selectively modulate mood states. This is also in line with our previous study showing that chronic but not acute activation of the whole ACC induces emotional deficits^10, 25^. Thus, it is likely that the development of a depressive-like phenotype following BLA-ACC activation requires long-term molecular and neuronal plasticity, which we investigated using open-ended approaches, and compared with human MDD signature.

Based on convergent bioinformatic analyses (common DEG, RRHO2, GSEA, WGCNA), we found that transcriptomic changes occurring in our optogenetic paradigm recapitulated a series of adaptations that were previously associated with human MDD, notably affecting the mitochondria^42, 59^, chromatin remodeling factors^60^, synaptic function, translational regulation^42^ or myelination^35, 61^. These results document the translational relevance of our paradigm. They also indicate that selective manipulation of a restricted neuronal pathway may represent a valuable strategy to model MDD, at both behavioral and molecular levels. Because recent studies suggest that the various mouse models available for this disorder may capture distinct aspects of its molecular pathophysiology^42^, we argue that our optogenetic model provides a complementary approach. While it is based on an artificially-induced neuronal hyperactivation, it appears to offer the possibility of modelling some of the effects of internal insults or states, such as chronic pain. Finally, this paradigm, and its potential extension to other neuronal circuits, also enables deciphering what is sufficient for the emergence of mood dysfunction.

Most widespread alterations affected oligodendrocytes and the myelination process, which were consistently identified across human and mouse data. Global down-regulation of myelin sheath proteins (*Plp1, Mal, Mog, Mbp*) and enzymes involved in the synthesis of myelin lipids (A*spa, Ugt8*), as well as up-regulation of myelination inhibitors (*Lingo1, Sema4a*) were observed in the ACC of animals displaying depressive-like behaviors. These results are congruent with previous studies reporting deficits in myelination^35, 62^, white matter tract organization^63–65^ or oligodendrocytes integrity^47, 66^, in the ACC of MDD patients. The crucial role of myelination in MDD pathophysiology is further supported by preclinical studies^49, 67, 68^. Depletion of the myelin sheath component CNP^50^, or of the positive regulator of myelination ErbB4^51^, as well as cellular depletion of oligodendrocyte progenitors^70^, have been shown to induce emotional dysfunction in rodents. Conversely, the pharmacological compound Clemastine, which enhances oligodendrocyte differentiation and myelination, exerts antidepressant-like effects in socially defeated,^68^ or isolated^66^ mice. However, molecular mechanisms mediating such effects remain poorly characterized. Here, using gene network theory, we prioritized *Sema4a* as one of the most prominently upregulated genes in a myelin-enriched gene module strongly affected by BLA-ACC activation. Previous reports have shown that increased *Sema4a* function contributes to the broad and profound loss of mature oligodendrocytes and demyelination observed in neurological and auto-immune disorders, such as multiple sclerosis^52–55, 72^. Our results extend these findings to a milder and more localized dysregulation of *Sema4a* signaling in the ACC, in the context of mood regulation. *Sema4a* knockdown in this region was sufficient to prevent the development of depressive-like behaviors induced by repeated activation of incoming BLA fibers. Therefore, we document a new role of *Sema4a*, whereby its optogenetically-driven increased ACC expression plays a pivotal role in mood dysregulation. Altogether, these results point to *Sema4a* as a new potential preclinical target for depression.

In conclusion, these results demonstrate that the BLA-ACC pathway has a selective but essential role at the crossroad of chronic pain and depression. By combining animal and human studies, we define the behavioral relevance of this pathway, and uncover the essential role of impaired myelination and *Sema4a* signaling in depression.

## Acknowledgements

This work was supported by the Centre National de la Recherche Scientifique (contract UPR3212), the University of Strasbourg, the Fondation pour la Recherche Médicale (FRM, FDT202012010622; FDT201805005527) (LJB, MH), the Fondation de France (FdF N° Engt:00081244; IY, PEL, RB, ECI), a NARSAD Young Investigator Grant from the Brain & Behavior Research Foundation (24736; IY), the French National Research Agency (ANR) through the Programme d’Investissement d’Avenir EURIDOL graduate school of pain ANR-17- EURE-0022 (LJB), ANR-18-CE37-0004 (IY), ANR-18-CE19-0006-03 (IY) and ANR-19-CE37-0010 (PEL), Hacettepe University Scientific Research Projects Coordination Unit (HUBAB), International Cooperation Project TBI-2018-17569 (BA), the Scientific and Technological Research Council of Turkey (TUBITAK) through international post-doctoral research fellowship program (BA), IdEx Young Investigator award (IY) and EU Erasmus Mundus Neurotime program (MK, IY, LH). The authors would also like to acknowledge the CAIUS High Performance Computing Center of the University of Strasbourg for providing scientific support and access to computing resources. Part of the computing resources was funded by the Equipex Equip@Meso project (Programme Investissements d’Avenir) and the CPER Alsacalcul/Big Data. We would like to thank Chronobiotron for animal care, Pascale Koebel and Paola Rossolillo from IGBMC for virus preparations, Jennifer Kaufling and Quentin Leboulleux for technical support, Violaine Alunni from IGBMC for the Fluidigm experiment and Daniel Almeida for English editing. Sequencing was performed by the GenomEast platform, a member of the ‘France Génomique’ consortium (ANR-10-INBS-0009).

## Authors contributions

Behavioral experiments: *LJB, RW, CF, SHJ, MH, IY*; Molecular experiments: *LJB, IY*; Electrophysiological recordings: *SH*; Human data acquisition: *GT, PEL*; Imaging: *LJB, MK, LD MM, MNS, LAH*; Immunohistochemistry: *RW, BA, MB*; Neuroanatomy: *RW, CF, PV*; Experimental design: *PEL, IY*; Data analyses: *LJB, RW, PH, BA, PEL, IY*; Fundings: *IY, PEL, ECI, RB*; Manuscript preparation: *LJB, RW, SH, LAH, PEL, IY*.

## Online methods

### Animals

*Experiments were conducted using male adult C57BL/6J (RRID: IMSR JAX: 000664) mice (Charles River), 8 weeks old at the beginning of experimental procedures, group-housed with a maximum of 5 animals per cage and kept under a reversed 12 h light/dark cycle. After the optic fiber implantation, animals were single housed to avoid possible damage to the implant. We conducted all the behavioral tests during the dark phase, under red light. Our animal facility (Chronobiotron) is registered for animal experimentation (Agreement A67-2018-38), and protocols were approved by the local ethical committee of the University of Strasbourg (CREMEAS, APAFIS8183-2016121317103584)*.

### Surgical procedures

*Surgical procedures were performed under zoletil/xylazine anesthesia (zoletil 50mg/ml, xylazine 2.5mg/ml; ip, 4ml/kg, Centravet). For stereotaxic surgery, a local anesthetic was delivered subcutaneously at incision site (bupivacaine, 2mg/kg)*.

### Neuropathic pain induction: cuff surgery

*For the BLA-ACC inhibition study, we used a well-characterized chronic pain-induced depression model*^11, 23^*. Before surgery, mice were assigned to experimental groups so that these groups did not initially differ in mechanical nociceptive threshold or body-weight. Chronic neuropathic pain was induced by placing a 2mm polyethylene tubing (Cuff, Harvard Apparatus, Les Ulis, France) around the right common branch of sciatic nerve*^24^*. The Sham group underwent the same procedure without cuff implantation*.

### Virus injection

*After general anesthesia, mice were placed in a stereotaxic frame (Kopf Instruments). 0.5µl of AAV5-CaMKIIa-ChR2(H134R)-EYFP or AAV5CaMKIIa-eArchT3.0-EYFP (UNC Vector core) were injected bilaterally into the BLA using a 5µl Hamilton syringe (0.05µl/min, coordinates for the BLA, anteroposterior (AP): -1.4mm from bregma, lateral (L): +/-3.2mm, dorsoventral (DV): 4mm from the brain surface). The same method was used to bilaterally inject the rAAV-mCherry-scrambleUsh or the rAAV-mCherry-Sema4Ash791 into the ACC (coordinates: AP: +0.7mm, L: +/-0.2mm, DV: -2mm, from the bregma). After injection, the 32-gauge needle remained in place for 10min before being removed and then the skin was sutured. Following surgery, animals were left undisturbed for at least two weeks before cannula implantation. To check viral injection localization at the end of the experiment, animals were anesthetized with Euthasol (182mg/kg) and perfused with 30mL of 0.1M phosphate buffer (PB, pH 7.4) followed by 100mL of 4% paraformaldehyde solution (PFA) in 0.1M PB. Brains were extracted and post fixed overnight and kept at 4°C in 0.1M PB saline (PBS) until cutting. Coronal sections (40µm) were obtained using a vibratome (VT 1000S, Leica, Deerfield, IL) and were serially collected in PBS. Sections were serially mounted with Vectashield medium (Vector laboratories) and localization of the fluorescence was checked using an epifluorescence microscope (Nikon 80i, FITC filter). Only animals well-injected bilaterally in the BLA were kept for further analyses*.

### Tracer injections

*Analysis of afferent neurons from the BLA to the ACC was performed by injecting the retrograde tracer Hydroxystilbamidine methanesulfonate (FluoroGold®,0.5µl) bilaterally in the ACC, using a microsyringe pump controller (UMC4, World precision instruments) and a 5µl Hamilton syringe (100nl/min, coordinates for the ACC, AP: +0.7mm from bregma, L: +/-0.2mm, DV: -2mm from the bregma). 7 days after the tracer injection, mice were anesthetized with Euthasol (182mg/kg) and perfused with 30ml of 0.1M phosphate buffer (PB, pH 7.4) followed by 150ml of 4% paraformaldehyde solution (PFA) in 0.1M PB. Brains were removed, postfixed overnight in PFA at 4°C, and then kept at 4°C in 0.1M PB saline (PBS, pH 7.4) until cutting. Coronal sections (40µm) were obtained with a Vibratome (VT 1000S, Leica, Deerfield, IL) and serially collected in PBS*.

### Optic fiber cannula implantation

*At least two weeks after virus injection, the animals underwent a unilateral optic fiber cannula implantation into the ACC. The optic fiber cannula was 1.7mm long and 220µm in diameter. The cannula was inserted 1.5mm deep in the brain at the following coordinates: AP: +0.7mm L: +/-0.2mm (MFC 220/250–0.66 1.7mm RM3 FLT, Doric Lenses)*^10^*. For behavioral experiments, cannulas were implanted in the left hemisphere in half of each experimental group, whereas the other half received the implant in the right hemisphere. For DTI protocol, all mice were implanted in the left hemisphere*.

### Optogenetic stimulation procedure

*After 3 to 7 days of recovery, the BLA-ACC pathway was activated or inhibited by a blue light emitting diode (LED) with a peak wavelength of 463nm (LEDFRJ-B FC, Doric Lenses) or a green light emitting laser with a peak wavelength of 520nm (Miniature Fiber Coupled Laser Diode Module, Doric Lenses) respectively. From the light source, the light travelled through the fiber optic patch cable (MFP 240/250/900-0.63 0.75m FC CM3, Doric Lenses) to the implant cannula. Blue light was delivered by pulses generated through a universal serial bus connected transistor-transistor logic pulse generator (OPTG 4, Doric Lenses) connected to a LED driver (LEDRV 2CH v.2, Doric Lenses). Transistor-transistor logic pulses were generated by an open-source software developed by Doric Lenses (USBTTL V1.9). Stimulated animals received repetitive stimulation sequences of 3s consisting of 2s at 10Hz with 10ms pulses and 1s without stimulation. The whole sequence is repeated during 20min each day for 3 consecutive days during 3 weeks. Light intensity was measured before implantation at the fiber tip using a photodetector (UNO, Gentec, Quebec, Canada) and was set between 3mW and 5mW. Green light was delivered in a continuous manner during 5min prior (Forced Swim Test and Dark/light test) or during behavioral testing (Splash test, Novelty Suppressed Feeding Test and von Frey test). The onset and end of stimulation were manually directed. Light intensity was measured as described above and set at 16mW. All control animals underwent the same procedure but the light remained switched off*.

### Behavioral assessment

#### Nociception-related behavior

*The mechanical threshold of hind paw withdrawal was evaluated using von Frey hairs (Bioseb, Chaville, France)*^23^*. Mice were placed in clear Plexiglas® boxes (7×9×7cm) on an elevated mesh screen*^24^*. Filaments were applied to the plantar surface of each hind paw in a series of ascending forces (0.4 to 8g). Each filament was tested five times per paw, being applied until it just bent, and the threshold was defined as 3 or more withdrawals observed out of the 5 trials. All animals were tested before the cuff surgery to determine the basal threshold, every week after cuff surgery to ensure the development of mechanical allodynia and during optogenetic stimulation to assess the effect of the inhibition of the BLA-ACC on mechanical hypersensitivity*.

#### Locomotor activity

*Spontaneous locomotor activity was monitored for each experimental group. Mice were individually placed in activity cages (32x20cm floor area, 15cm high) with 7 photocell beams. The number of beam breaks was recorded over 30min using Polyplace software (Imetronic, Pessac, France)*.

#### Real time place avoidance (RTA) and Conditioned Place Preference (CPP)

*The apparatus consists of 2 connected Plexiglas chambers (size 20cmx20cmx30 cm) distinguished by the wall patterns. On the first day (pre-test), animals are free to explore the apparatus during 5min (CPP) or 10min (RTA), and the time spent in each chamber is measured to control for the lack of spontaneous preference for one compartment. Animals spending more than 75% or less than 25% of the total time in one chamber were excluded from the study. For RTA, the second day (test), animals are plugged to the light source placed between the 2 chambers and let free to explore for 10min. Light is turned on when the mouse enters its head and forepaws in the stimulation paired chamber and turned off when it quit the compartment. Total time spent in the stimulation-paired chamber is measured. For CPP, on the second and third days (conditioning), animals are maintained during 5min in one chamber, where optogenetic stimulation occurs, and 4h later placed during 5min in other chamber, without optogenetic stimulation. On the 4th day (test), the time spent in each chamber is recorded during 5min*.

### Anxiodepressive-like behavioral assays

*Behavioral testing was performed during the dark phase, under red light. While each mouse went through different tests, they were never submitted to more than 2 tests per week, and never went through the same test twice. The forced swimming test (FST) was always considered as terminal. Body weights were measured weekly*.

#### Dark-Light Box Test

*The apparatus consisted of light and dark boxes (18×18×14.5 cm each). The lit compartment was brightly illuminated (1000lux). This test evaluates the conflict between the exploratory behavior of the rodent and the aversion created by bright light. Mice were placed in the dark compartment in the beginning of the test, and the time spent in the lit compartment was recorded during 5min*^10^*. For inhibition experiment, the test was performed immediately after the light stimulation*.

#### Novelty suppressed feeding test

*The apparatus consisted of a 40×40×30 cm plastic box with the floor covered with 2cm of sawdust. Twenty-four hours prior to the test, food was removed from the home cage. At the time of testing, a single pellet of food was placed on a paper in the center of the box. The animal was then placed in a corner of the box and the latency to eat the pellet was recorded within a 5min period. This test induces a conflict between the drive to eat the pellet and the fear of venturing in the center of the box*^73^*. For inhibition experiments, optogenetic stimulation was conducted during the test*.

#### Splash Test

*This test, based on grooming behavior, was performed as previously described*^23, 73^*. Grooming duration was measured during 5min after spraying a 20% sucrose solution on the dorsal coat of the mouse. Grooming is an important aspect of rodent behavior and decreased grooming in this test is considered related to the loss of interest in performing self-oriented minor tasks*^74^*. For inhibition experiments, optogenetic stimulation was conducted during the test*.

#### Nest Test

*This test, based on a rodent innate behavior, was performed in cages identical to the home cages of animals. Each mouse was placed in a new cage with cotton square in the center. Water and food were provided ad libitum. After 5h, mice were placed back in their original cages and pictures of the constructed nest were taken. A score was given blindly to each nest as follows: 0 corresponds to an untouched cotton square, 1 to a cotton square partially shredded, 2 if the cotton is totally shredded but not organized, 3 if cotton is totally shredded and organized in the center of the cage, 4 if the cotton is totally shredded and shows a well-organized shape in the corner of the cage, like a nest*^75, 76^.

#### Forced Swim Test

*FST*^77^ *was conducted by gently lowering the mouse into a glass cylinder (height 17.5cm, diameter 12.5cm) containing 11.5cm of water (23-25°C). Test duration was 6min. The mouse was considered immobile when it floated in the water, in an upright position, and made only small movements to keep its head above water. Since little immobility was observed during the first 2min, the duration of immobility was quantified over the last 4min of the 6min test. Concerning inhibition experiments, the test was performed just after the stimulation*.

### Ex vivo electrophysiological recordings

*We performed whole-cell patch clamp recordings of BLA neurons or ACC pyramidal neurons. In the BLA, we recorded from eYFP-expressing neurons of mice bilaterally injected with an AAV driving the expression of either the archaerhodopsin ArchT3.0 or the channelrhodopsin 2 under control of the CaMKIIa promoter (with AAV5-CamKIIa-ArchT3.0-EYFP and AAV5-CaMKIIa-ChR2(H134R)-EYFP, respectively). In the ACC, we recorded from pyramidal neurons surrounded by eYFP-positive fibres. For these experiments, mice were anaesthetized with urethane (1.9g/kg) and killed by decapitation, their brain was removed and immediately immersed in cold (0°C-4°C) sucrose-based ACSF containing the following (in mm): 2 kynurenic acid, 248 sucrose, 11 glucose, 26 NaHCO_3_, 2 KCl, 1.25 KH_2_PO_4_, 2 CaCl_2_, and 1.3 MgSO_4_ (bubbled with 95% O_2_ and 5% CO_2_). Transverse slices (300μm thick) were cut with a vibratome (VT1000S, Leica). Slices were maintained at room temperature in a chamber filled with ACSF containing the following (in mM): 126 NaCl, 26 NaHCO_3_, 2.5 KCl, 1.25 NaH_2_PO_4_, 2 CaCl_2_, 2 MgCl_2_, and 10 glucose (bubbled with 95% O_2_ and 5% CO_2_; pH 7.3; 310mOsm measured). Slices were transferred to a recording chamber and continuously superfused with ACSF saturated with 95% O_2_ and 5% CO_2_. BLA neurons expressing eYFP were recorded in the whole-cell patch-clamp configuration. Recording electrodes (3.5-4.5MΩ) were pulled from borosilicate glass capillaries (1.2mm inner diameter, 1.69mm outer diameter, Warner Instruments, Harvard Apparatus) using a P1000 electrode puller (Sutter Instruments). Recording electrodes were filled with, in mM: 140 KCl, 2 MgCl_2_, 10 HEPES, 2 MgATP; pH 7.3. The pH of intrapipette solutions was adjusted to 7.3 with KOH, and osmolarity to 310mOsm with sucrose. BLA or ACC were illuminated with the same system used for the in vivo experiments (see above) triggered with WinWCP 4.3.5, the optic fiber being localized in the recording chamber at 3mm from the recorded neuron. The holding potential was fixed at -60mV. Recordings were acquired with WinWCP 4.3.5 (courtesy of Dr. J. Dempster, University of Strathclyde, Glasgow, United Kingdom). All recordings were performed at 34°C*.

### MRI data acquisition

*Mouse brain resting state functional MRI scans were performed with a 7T Bruker BioSpec 70/30 USR animal scanner, a mouse head adapted room temperature surface coil combined with a volume transmission coil for the acquisition of the MRI signal and ParaVision software version 6.0.1 (Bruker, Ettlingen, Germany). Imaging was performed at baseline, 2 weeks and 8 weeks after peripheral nerve injury in the cuff model (Cuff n=7 and Sham n=7). For rsfMRI the animals were briefly anesthetized with isoflurane for initial animal handling. The anesthesia was further switched to medetomidine sedation (MD, Domitor, Pfizer, Karlsruhe, Germany), initially induced by a subcutaneous (sc) bolus injection (0.15mg MD per kg body weight (kg bw) in 100μl 0.9% NaCl-solution). 10min later, the animals received a continuous sc infusion of MD through an MRI compatible catheter (0.3mg/kg bw/h) inserted at the mouse shoulder level. During the whole acquisition a 2mm thick agar gel (2% in NaCl) was applied on the mouse head to reduce any susceptibility artifacts arising at the coil/tissue interface. Respiration and body temperature were monitored throughout the imaging session. Acquisition parameters for rs-fMRI were: single shot GE-EPI sequence, 31 axial slices of 0.5mm thickness, FOV=2.12×1.8cm, matrix=147×59, TE/TR=15ms/2000ms, 500 image volumes, 0.14×0.23×0.5mm^3^ resolution. Acquisition time was 16min. Morphological T2-weighted brain images (resolution of 0.08×0.08×0.4mm^3^) were acquired with a RARE sequence using the following parameters: TE/TR=40ms/4591ms; 48 slices, 0.4mm slice thickness, interlaced sampling, RARE factor of 8, 4 averages; an acquisition matrix of 256×256 and FOV of 2.12×2cm^2^. Brain Diffusion Tensor MRI (DT-MRI) acquisition in the BLA-ACC optogenetically stimulated animals was performed with the 7T animal scanner, but using a combination of a transmit – receive volume coil (86mm) and a mouse brain adapted loop surface coil allowing the passage of the optogenetic cannulas (MRI, Bruker, Germany). Stimulated animals (n=6) and their controls (n=7) were brain imaged under isoflurane anesthesia (1.5% for maintenance) using a 4-shot DTI-EPI sequence (TE/TR=24ms/3000ms), 8 averages; with diffusion gradients applied along 45 nonlinear directions, gradient duration [δ]=5.6ms and gradient separation [Δ]=11.3ms and a b-factor of 1000s/mm^2^. Images with a b-factor=0s/mm2 were also acquired. 30 axial slices with 0.5mm thickness were acquired covering the whole brain with a FOV of 1.9×1.6cm^2^ and an acquisition matrix of 190×160 resulting in an image resolution of 0.1×0.1×0.5mm^3^. The total acquisition time was of 1h20min*.

### MRI data processing

*Rs-fMRI images were spatially normalized into a template using Advanced Normalization Tools (ANTs) software*^78^ *using SyN algorithm and smoothed (FWHM=0.28×0.46×1mm^3^) with SPM8. Seed-based functional connectivity analysis was performed with a MATLAB tool developed in-house. Regions of interest (ROI) were extracted from Allen Mouse Brain Atlas*^37^ *which were later normalized into the template space. Resting-state time series were de-trended, band-pass filtered (0.01-0.1Hz) and regressed for cerebrospinal fluid signal from the ventricles. Principal component analysis (PCA) of the BOLD time courses across voxels within a given ROI was performed and first principal component accounting for the largest variability was selected as the representative time course for further analysis. Spearman correlations between the PCA time course of single ROIs and each voxel of the brain was computed at the group and individual levels and r values were converted to z using Fisher’s r-to-z transformation. Individual connectivity maps for baseline rs-fMRI acquisitions were subtracted from 2 and 8 PO weeks counterparts for each subject. Baseline subtracted connectivity maps were subsequently used for two sample t-test with SPM8 to perform group comparison. Family-wise error rate (FWER) correction was applied at the cluster level (p<0.05) for each statistical image. Preprocessing of diffusion weighted images included denoising*^79^*, removal of Gibbs ringing artifacts*^80^*, motion correction*^81^*, and bias field inhomogeneity correction*^82^*. Diffusion tensor was estimated*^83^ *using weighted least-squares (WLS) approach and the following tensor-derived parameters were computed: fractional anisotropy (FA), axial diffusivity (AD), mean diffusivity (MD) and radial diffusivity (RD). All these processing steps were done using MRtrix3 (*https://www.mrtrix.org*); except the motion correction step, done using Advanced Normalisation Tools (ANTs,* http://stnava.github.io/ANTs/*). Based on b=0s/mm*^2^ *images (i.e., the volume without diffusion weighting), these images were then spatially registered in a common space using the SyN registration method of ANTs to build a study-specific template, which was then affinely registered onto the Allen Brain Atlas template. Each mouse tensor-derived maps were finally warped in this common space. These registered images were finally smoothed by a 0.5mm full-width half maximum (FWHM) Gaussian kernel. Inter-group differences for all the DTI derived parameters were assessed using the SPM12 General Linear Model (GLM). The results were analyzed according to a level of statistical significance, p<0.001 without correction. Further, correlational analyses were performed between the DTI metrics (voxel level) and the results from splash tests (statistical significance was p<0.001, uncorrected)*.

### Immunohistochemistry

#### c-Fos immunoperoxydase

*Animals were stimulated once with the same procedure as described before (for BLA-ACC activation). 90min later, animals were anesthetized with Euthasol (182mg/kg) and perfused with 30ml of 0.1M PB (pH 7.4) followed by 100ml of 4% PFA in 0.1M PB. Brains were removed, post fixed overnight and kept at 4°C in 0.1M PBS (pH 7.4) until cutting. Coronal sections (40µm) were obtained using a vibratome (VT 1000S, Leica, Deerfield, IL) and were serially collected in PBS. Sections were incubated 15min in a 1% H_2_O_2_/50% ethanol solution and washed in PBS (3×10 min). Sections were then pre-incubated in PBS containing Triton X-100 (0.3%) and donkey serum (5%) for 45min. Sections were then incubated overnight at room temperature in PBS containing Triton X-100 (0.3%), donkey serum (1%) and rabbit anti-c-Fos (1:10000, Santa Cruz Biotechnology, E1008). Sections were then washed in PBS (3×10min), incubated with biotinylated donkey anti-rabbit secondary antibody (1:300) in PBS containing Triton X-100 (0.3%), donkey serum (1%) for 2h and washed in PBS (3×10min). Sections were incubated with PBS containing the avidin-biotin-peroxidase complex (ABC kit; 0.2% A and 0.2% B; Vector laboratories) for 90min. After being washed in Tris-HCl buffer, sections were incubated in 3,3’diaminobenzidine tetrahydrochloride (DAB) and H_2_O_2_ in Tris-HCl for approximately 4min and washed again. Sections were serially mounted on gelatin-coated slides, air dried, dehydrated in graded alcohols, cleared in Roti-Histol (Carl Roth, Karlsruhe, Germany) and coverslipped with Eukitt. c-Fos immunohistochemistry then allowed controlling for both the implant location and the activation of the ACC by the optogenetic procedure. Animals having c-Fos induction outside of the ACC, for instance in the motor cortex, were excluded from analysis*.

#### c-Fos immunofluorescence

*Animals were anesthetized with Euthasol (182mg/kg) and perfused with 30ml of 0.1M PB (pH 7.4) followed by 100ml of 4% PFA in 0.1M PB. Brains were removed, postfixed overnight and kept at 4°C in 0.1M PBS (pH 7.4) until cutting. Coronal sections (40µm) were obtained using a vibratome (VT 1000S) and were serially collected in PBS. Sections were washed in PBS (3×10min) and pre-incubated in PBS containing Triton X-100 (0.3%) and donkey serum (5%) for 45min. Sections were then incubated overnight at room temperature in PBS containing Triton X-100 (0.3%), donkey serum (1%) and rabbit anti-c-Fos (1:1000, Synaptic System, 226-003). Sections were then washed in PBS (3×10min), incubated with Alexa fluor 594 donkey anti-rabbit secondary antibody (1:400) in PBS containing Triton X-100 (0.3%), donkey serum (1%) for 2h and washed in PBS (3×10min). Sections were finally serially mounted with vectashield medium (Vector laboratories)*.

### Fluorogold and cFos quantification

*Single-layer images were acquired using a laser-scanning microscope (confocal Leica SP5 Leica Microsystems CMS GmbH) equipped with x20 objective. Excitation wavelengths were sequentially diode 405nm, argon laser 488nm and diode 561nm. Emission bandwidths are 550-665nm for Fluorogold fluorescence and 710-760nm for Alexa594 signal. Segmentation and classification of c-Fos positive cells was performed from 3 sections for each animal using a deep learning model. The model was trained from scratch for 400 epochs on 10 paired image patches (image dimensions: (160,160), patch size: (160,160)) with a batch size of 2 and a mae loss function, using the StarDist 2D ZeroCostDL4Micnotebook (v 1.11)*^84^*. Key python packages used include tensorflow (v 0.1.12), Keras (v 2.3.1), csbdeep (v 0.6.2), numpy (v 1.19.5), cuda (v 11.0.221). The training was accelerated using a Tesla K80 GPU and dataset was argumented by a factor of 4. Segmentation and classification of Fluorogold signal was done using Stardist 2D_versatile_fluo*^85, 86^ *pre-trained model. Fluorogold masks and c-fos masks were then overlaid to count the double positive cells using Fiji*^87^.

### RNAscope

*Brain samples were immersed in isopentane and immediately placed at -80°C. Frozen samples were embedded in OCT compound and 14µm thick sections were performed on cryostat, mounted on slides and put back in -80°C freezer. Sections were fixed, dehydrated and pre-treated using the “RNAscope Sample Preparation and Pre-treatment Guide for Fresh Frozen Tissue using RNAscope Fluorescent Multiplex Assay” protocol (Advanced Cell Diagnostics). Hybridation of Slc17a7 (ACD, 416631), Gad2 (ACD, 415071-C2) and c-fos (ACD, 316921) probes and development of the different signals with Opal 520, 590 and 690 fluorophores were performed in accordance with the “RNAscope Multiplex Fluorescent Reagent Kit v2 Assay” instructions (Advanced Cell Diagnostics). Single-layer images were acquired using a laser-scanning nanozoomer (S60; Hamamatsu Photonics) at 40X magnification. Quantifications were performed from two sections for each animal on QuPath 0.3.0 software*^88^*. First, region of interest were drawn using the polygon annotation tool. Then nuclei were detected within regions of interest using the cell detection module on the Dapi staining. To determine the c-fos, Gad2 and Slc17a7 positive cells on our regions of interest, object classifiers were trained in Qupath using Random trees classifiers. We selected all the features by output class (Nucleus mean, Nucleus sum, Nucleus standard deviation, Nucleus maximum, Nucleus minimum, Nucleus range, Cell mean, Cell standard deviation, Cell maximum, Cell minimum, Cytoplasm mean, Cytoplasm standard deviation, Cytoplasm maximum, Cytoplasm minimum), and annotated manually a minimum of 20 points for positive cells and negative cells. Classifiers were then applied sequentially on the whole region of interest to determine the c-fos, Gad2 and Slc17a7 positive cells*.

### RNA extraction

*Two different batches of animals were generated for RNA-sequencing, with a third one for Fluidigm validation of RNA-sequencing results. Bilateral ACC was freshly dissected from animals killed by cervical dislocation and tissues were stored at -80°C. Total RNA was extracted from ACC tissue with the Qiagen RNeasy Mini Kit (Hilden Germany). Around 20mg of ACC tissue was disrupted and homogenized with a Kinematica Polytron 1600E in 1.2ml QIAzol Lysis reagent, for 30s, and then left at room temperature for 5min. Next, 240µl of chloroform was added and mixed before centrifugation for 15min at 12,000rpm at 4°C. The aqueous phase (600µl) was transferred to a new collection tube and mixed with 600µl of 70% ethanol. The mix was transferred into a RNeasy spin column in a 2ml collection tube, and centrifuged at 10,000rpm for 15s. Next, 350µl of RW1 buffer was added and centrifuged at 10,000rpm, for 15s, before adding 10µl of DNAse and 70µl of RDD buffer. The mix was left at room temperature for 15min and 350µl of RW1 buffer was added and centrifuged at 10,000rpm for 15s. The column was then transferred to a new 2ml collection tube and washed with 500µl of RPE buffer, before being centrifuged at 10,000rpm. Finally, the column was dry centrifuged at 10,000rpm for 5min, and transferred to a new 1.5ml collection tube to which 18µl of RNase-free water was added. Finally, the RNA was eluted by centrifugation for 1min at 10,000rpm. Samples were kept at -80°C until use*.

### Mouse RNA-sequencing

*RNA-sequencing was performed by the Genomeast platform at IGBMC. Full length cDNAs were generated from 5ng of total RNA using the Clontech SMARTSeq v4 Ultra Low Input RNA kit for Sequencing (PN 091817, Takara Bio Europe, Saint-Germain-en-Laye, France) according to manufacturer’s instructions, with 10 cycles of PCR for cDNA amplification by Seq-Amp polymerase. Six hundred pg of pre-amplified cDNA were then used as input for Tn5 transposon tagmentation by the Nextera XT DNA Library Preparation Kit (PN 15031942, Illumina, San Diego, CA), followed by 12 PCR cycles of library amplification. Following purification with Agencourt AMPure XP beads (BeckmanCoulter, Villepinte, France), the size and concentration of libraries were assessed by capillary electrophoresis. Libraries were then sequenced using an Illumina HiSeq 4000 system using single-end 50bp reads. Reads were mapped onto the mm10 assembly of the Mus musculus genome, using STAR version 2.5.3a*^89^*. Gene expression quantification was performed from uniquely aligned reads using htseq-count*^90^*version 0.6.1p1, with annotations from Ensembl version 95. Read counts were then normalized across samples with the median-of-ratios method proposed by Anders and Huber*^91^*, to make these counts comparable between samples. Principal Component Analysis was computed on regularized logarithm transformed data calculated with the method proposed by Love and collaborators*^92^*. Differential expression analysis was performed using R and the Bioconductor package DESeq2 version 1.22.1*^92^*, using RIN values and batches as covariates. Because we generated two batches of mice, the lfcShrink function was used instead of betaPrior in order to calculate p-values from the log2 Fold-changes unshrinked and to perform the shrinkage afterwards. RIN and sample batches can be found in **Extended Data Table2***.

### Human RNA-Sequencing data

*Human gene expression data, obtained from our previous publication*^35^ *(archived on GEO Datasets under the reference series: “GSE151827” samples: GSM5026548-97), were generated initially using post-mortem ACC tissue from the Douglas-Bell Canada Brain Bank. This cohort was composed of 26 subjects who died by suicide during a major depressive episode, and 24 psychiatrically healthy controls. Groups were matched for age, post-mortem interval and brain pH, and include both male (19 in control group and 19 in MDD group) and female (5 in control group, 7 in MDD group) subjects*. *Demographics for the cohort can be found in **Extended Data Table2**. While differential expression analysis for the whole cohort (both males and females) was reported previously*^35^*, during the present work we reprocessed raw gene counts from male individuals only, and conducted a new differential expression analysis to compare men with MDD and men healthy control, taking into account RIN, and age, as in*^35^.

### Rank-rank hypergeometric overlap (RRHO) analysis

*In order to compare mouse and human RNA-Sequencing data, we used the Rank-rank hypergeometric overlap (RRHO2) procedure, as described by Cahill et al*^36^*, using the R package available at:* https://github.com/Caleb-Huo/RRHO2*. Mice-human orthologous genes were first obtained using the R package BioMart, leaving a total of 13572 genes. Genes in each data set were ranked based on the following metric: -log10(p-value) x sign(log2 Fold Change). Then, the RRHO2 function was applied to the 2 gene lists at default parameters (with stepsize equal to the square root of the list length). Significance of hypergeometric overlaps between human and mouse gene expression changes are reported as log10 p-values, corrected using the Benjamini–Yekutieli procedure*.

### Gene Set Enrichment Analysis (GSEA)

*Mouse and men genes were ranked independently based on the fold changes obtained from their respective differential expression analysis. GSEA was performed as previously described*^93^ *using the he GSEAPreranked tool and the Lein Oligodendrocyte markers gene set*.

### Weighted Gene Coexpression Network Analysis (WGCNA)

*WGCNA*^38^ *was used to construct gene networks in mice and human using RNA-seq expression data and then identify conserved gene modules between the two species. The RNA-sequencing expression data were normalized for batch and RIN in mice; and for age, ethnic origin and RIN in human (with sex included when analyzing the whole cohort). First, a soft-threshold power was defined (mouse: 4, human: 8) to reach a degree of independence superior to 0.8 and thus ensure the scale-free topology of the network. To construct the network and detect modules, the blockwiseModules function of the WGCNA algorithm was used, with the minimum size of modules set at 30 genes. Then, the eigengene of each module was correlated with our traits of interest (optogenetic stimulation of the BLA-ACC pathway in mice, or MDD in human) and gene significance (GS), defined as the correlation between each individual gene and trait, was calculated. Inside each module a measure of the correlation between the module eigengene and the gene expression profile, or module membership (MM), was also assessed. Conservation of WGCNA module across mice and human was assessed by Fisher’s exact test. Modules were considered as significantly overlapping, and therefore conserved, when padj<0.05. Among the modules displaying a significant overlap between human and mice, only those with a significant (padj<0.1) association between the module eigengene and trait, in both species, were kept for further analysis*.

### Gene Ontology

*Enrichment for functional terms in DEGs in human MDD and mice was performed using WEBGSTALT for biological process, cellular component and molecular function*^34^*. Analysis was restricted to the genes differentially expressed at nom-p<0.05. The same procedure was applied to the list of genes changed in the same direction in mice and human obtained by RRHO without regards on the p-value*.

### Fluidigm

*cDNA was generated by subjecting 50ng of RNA from each sample to reverse transcriptase reaction (Reverse Transcription Master Mix Kit Fluidigm P/N-100-6297). Then, 1.25µL of each cDNA solution was used to generate a preamp mix containing a pooled of the 26 primers pairs and the PreAmp Master Mix Kit (Fluidigm P/N 100-5744). Preamp mixes were run for 14 cycles and the remaining primers were digested with Exonuclease I (New England BIOLAB. P/N M0293l. LOT 0191410). Preamp samples were analyzed for the expression of 22 genes of interest (for primer sequences see **Extended data TableS8**) using the BioMark qPCR platform (Fluidigm, San Francisco, CA, USA). Data were normalized to Gadph, B2m, Actb and Gusb (from the same animal) and fold changes were calculated using the 2-ΔΔCt method*^94^.

### Plasmid construction

*For the expression of shRNAs, long oligonucleotide linkers were designed containing the HindIII and BglII restriction sites at the 5’ and 3’ extremities respectively. Each linker contained the loop TTCAAGAGA separating a forward and a reverse copy of the following shRNA sequences: GGAAGAGCCAGACAGGTTTCT for mouse Sema4A exon15, GCCACAACGTCTATATCATGG for eGFP and GCGCTTAGCTGTAGGATTC for a universal scramble. Linkers were cloned by restriction/ligation downstream the mouse U6 promoter into the pAAV-CMV-mCherry-mU6 construct derived from pAAV-MCS (Agilent). For the construction of the plasmid for the sh efficiency testing in cells, the Sema4A exon 15 was amplified from C57Bl/6 embryonic stem cell genomic DNA and cloned at the C-terminal end of eGFP into pEGFP-C1 by SLIC using the following primers: GTACAAGTCCGGACTCAGATCTCGAGCTATTAAAGAAGTCCTGACAGTCCC and GATCAGTTATCTAGATCCGGTGGATCCTTAAGCCACTTCGGCGCC*.

### AAV production

*Recombinant adeno-associated virus AAV serotype 5 (AAV5) were generated by a triple transfection of HEK293T-derived cell line using Polyethylenimine (PEI) transfection reagent and the 3 following plasmids: pAAV-CMV-mCherry-mU6-shRNA, pXR5 (deposited by Dr Samulski, UNC Vector Core) encoding the AAV serotype 5 capsid and pHelper (Agilent) encoding the adenovirus helper functions. 48h after transfection, AAV5 vectors were harvested from cell lysate treated with Benzonase (Merck) at 120U/mL. They were further purified by gradient ultracentrifugation with Iodixanol (OptiprepTM density gradient medium) followed by dialysis and concentration against Dulbecco’s PBS using centrifugal filters (Amicon Ultra-15 Centrifugal Filter Devices 100K, Millipore). Viral titers were quantified by Real-Time PCR using the LightCycler480 SYBR Green I Master (Roche) and primers targeting mCherry sequence. Titers are expressed as genome copy per milliliter (GC/mL)*.

### Statistical analyses

*Statistical analyses were performed in GraphPad Prism v9.0 software. Data are expressed as mean±SEM, with statistical significance set as *p<0.05, **p<0.01, ***p<0.001. Student’s t-test (paired and unpaired), One-Way Analysis of Variance (ANOVA), One-Way Repeated Measures ANOVA, and Two-way ANOVA followed by Newman-Keuls post hoc test were used when appropriate. If data failed the Shapiro-Wilk normality test, Mann-Whitney non-parametric (one- or two-tailed) analysis was used*.

