## Supplementary Figures and Tables for "The basolateral amygdala-anterior cingulate pathway contributes to depression and its comorbidity with chronic pain"

### Description of the Extended data

Supplementary Figures 1-6, Supplementary Table 2, 7 and 8 are regrouped in **Extended data file**. Other Supplementary Tables (1 and 3 to 6) are displayed separately.

**Supplementary Table 1**:
Mouse differential expression analysis, gene ontology

**Supplementary Table 2**:
Human and mouse cohort metrics

**Supplementary Table 3**:
Gene Ontology of mice and men overlap computed by RRHO2

**Supplementary Table 4**:
Gene Ontology of mice and human (women + men) overlap computed by RRHO2

**Supplementary Table 5**:
Fisher table for the overlap between mice and men modules computed by WGCNA

**Supplementary Table 6**:
Fisher table for the overlap between mice and human (women + men) modules computed by WGCNA

**Supplementary Table 7**:
GO term enriched in the 5 mouse modules significantly associated with optogenetic stimulation and conserved in human

**Supplementary Table 8**:
Primers sequences for RT-qPCR (Fluidigm)

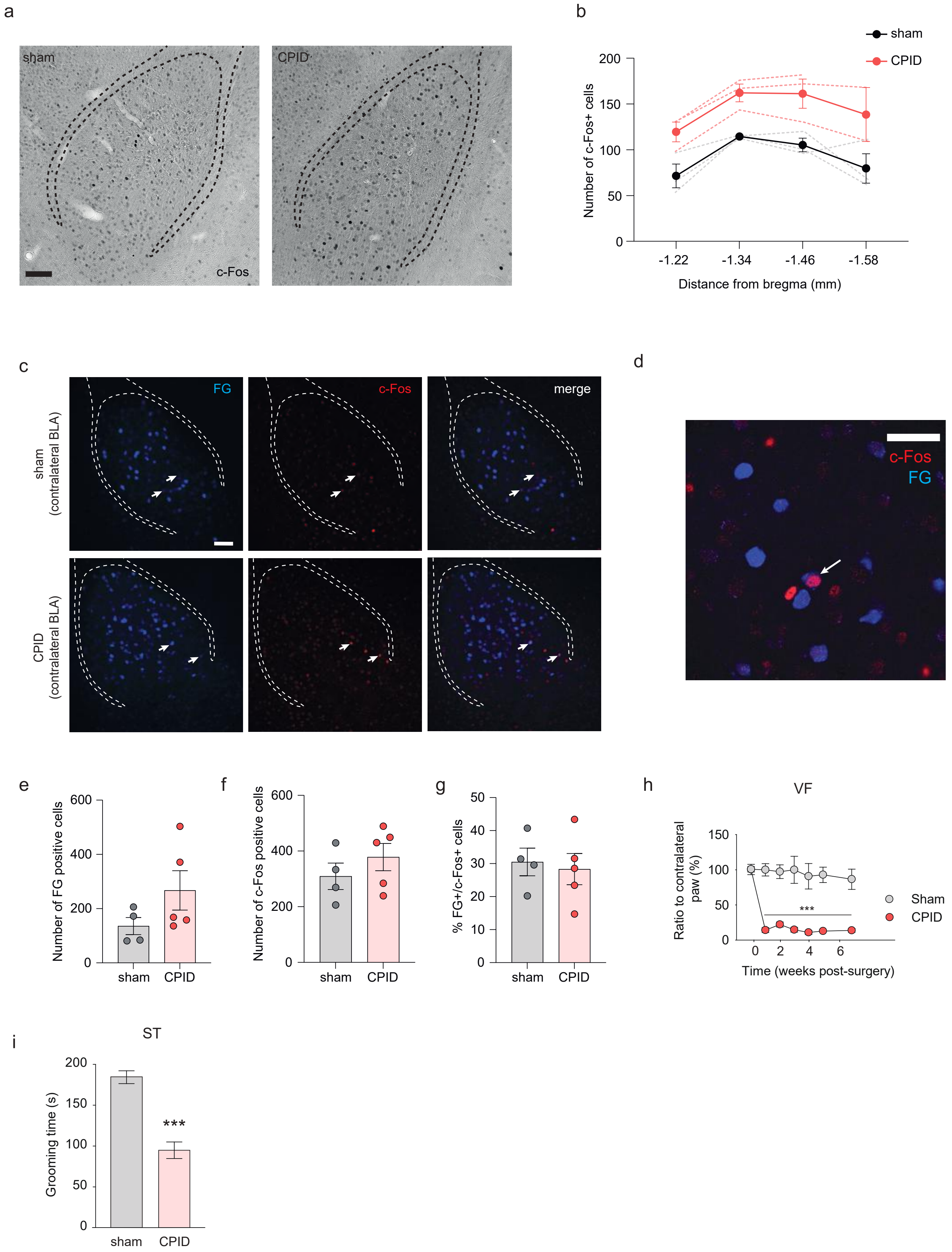
a
sham
CPID
c-Fos
b
sham
CPID
Number of c-Fos+ cells
200
150
100
50
0
-1.22
-1.34
-1.46
-1.58
Distance from bregma (mm)
c
FG
c-Fos
merge
sham (contralateral BLA)
CPID (contralateral BLA)
d
c-Fos
FG
e
Number of FG positive cells
600
400
200
0
sham
CPID
f
Number of c-Fos positive cells
600
400
200
0
sham
CPID
g
% FG+/c-Fos+ cells
50
40
30
20
10
0
sham
CPID
h
VF
Ratio to contralateral paw (%)
150
100
50
0
***
0
2
4
6
Time (weeks post-surgery)
Sham
CPID
i
ST
Grooming time (s)
200
150
100
50
0
***
sham
CPID

**Figure S1**. **a**. Representative images showing c-Fos immunoreactivity in the left (contralateral to the nerve injury) BLA of sham (left panel) and CPID (right panel) animals at 8 weeks PO. Scale bars=100µm **b**. The number of c-Fos positive cells was increased in the BLA in CPID animals (ctrl: n=3; stim: n=3; $F_{(1,4)}$=15.44; p=0.017, dotted traces=individual responses). **c**. Representative fluorescence images showing Fluorogold positive (FG+, left panel), c-Fos positive (c-Fos+, middle panel) cells and their co-localization (right panel) in the left (contralateral to the nerve injury) BLA after FG injection into the ACC. Scale bar=100µm. **d.** Close-up image showing c-Fos and FG positive cell co-localization. Scale bar=20µm. **e-g**. Quantification of FG+, c-Fos+ cells and their co-localization revealed that 8 weeks after cuff surgery, the number of FG+ (d; sham: 135.8±31.52; CPID: 267.2±72.58; p=0.21), c-Fos+ (e; sham: 309.3±47.60; CPID: 378.2±49.10; p=0.14) and FG+/c-Fos+ cells (f; sham 30.50±4.20; CPID: 28.33±4.72; p=0.45) were not altered in the left BLA (contralateral to the nerve injury; sham: n=4; CPID: n=5). **h-i.** Peripheral nerve injury induced an ipsilateral long-lasting mechanical hypersensitivity (h; sham: n=7; cuff: n=7; $F_{(6,72)}$=6.2629; p<0.0001; post-hoc weeks 1-7 p<0.05) and decreases grooming behavior in the splash test (i; sham: n=7; 185.21±19.07; CPID: n=7; 95.21±27.11; p=0.00001). Data are represented as mean ± SEM. 2-Way ANOVA (anterioposteriority x Surgery: c-Fos quantification); 2-Way ANOVA repeated measures (Time x Surgery; VF); one-tailed Mann-Whitney test (FG, c-Fos quantification). PO, post-operative; ST, Splash test; VF, von Frey filament test.

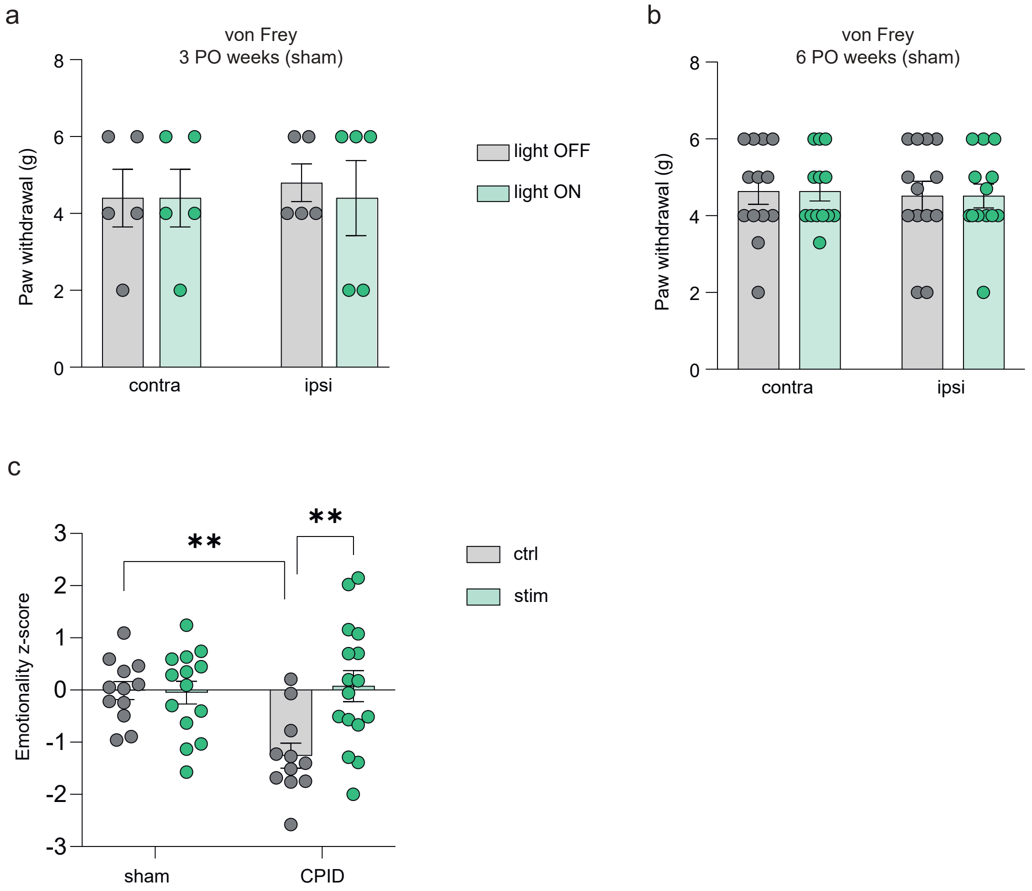

a
von Frey
3 PO weeks (sham)
Paw withdrawal (g)
0
2
4
6
8
contra
ipsi
light OFF
light ON
b
von Frey
6 PO weeks (sham)
Paw withdrawal (g)
0
2
4
6
8
contra
ipsi
c
Emotionality z-score
3
2
1
0
-1
-2
-3
**
**
sham
CPID
ctrl
stim

**Figure S2**. At 3 (**a**) or 6 (**b**) weeks after peripheral nerve injury, mechanical threshold was not altered by optogenetic inhibition of the BLA-ACC pathway in sham animals (ipsilateral vs contralateral paw; 3 PO weeks $F_{(1,4)}$=0.12; p=0.75; 6 PO weeks $F_{(1,12)}$=0.19; p=0.67; light-off vs light-on; 3 PO weeks $F_{(1,4)}$=0.12; p=0.75; 6 PO weeks $F_{(1,12)}$=0.00; p>0.99). **c**. CPID animals had decreased emotionality z-score which were reversed by optogenetic inhibition of the BLA-ACC pathway ($F_{(1,49)}$=7.488; p=0.008; post-hoc: sham-ctrl (n=12)>CPID-ctrl (n=11); p<0.01; CPID-ctrl (n=11)<CPID-stim (n=16); p<0.01; sham-ctrl (n=12)=sham-stim (n=14); p>0.05). Data are represented as mean±SEM. **p<0.01. Two-way ANOVA repeated measures (von Frey), two-way ANOVA (surgery x stimulation; Emotionality z-score).

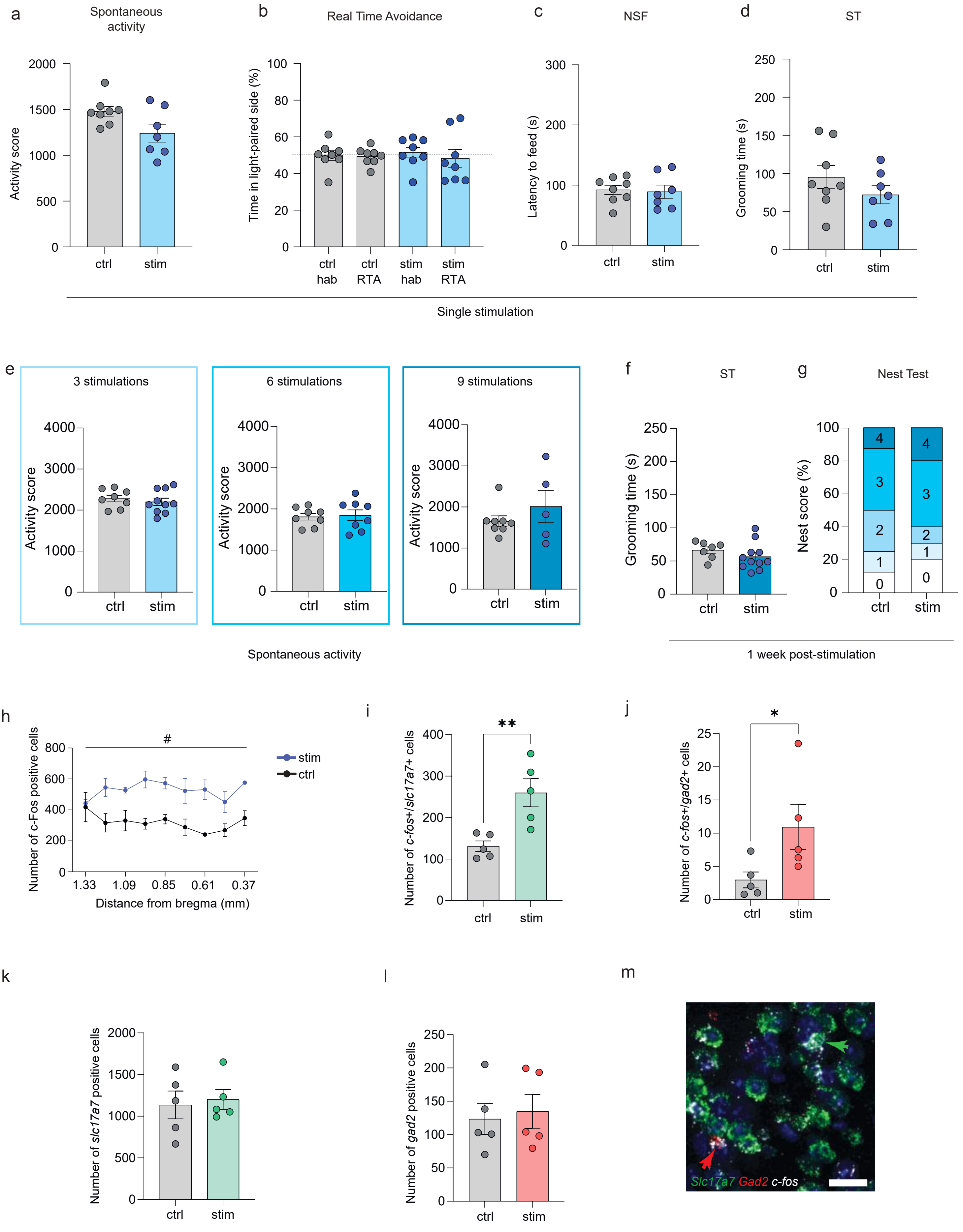

a
Spontaneous activity
Activity score
ctrl
stim
b
Real Time Avoidance
Time in light-paired side (%)
ctrl hab
ctrl RTA
stim hab
stim RTA
c
NSF
Latency to feed (s)
d
ST
Grooming time (s)
Single stimulation
e
3 stimulations
6 stimulations
9 stimulations
Spontaneous activity
f
ST
g
Nest Test
Nest score (%)
1 week post-stimulation
h
Number of c-Fos positive cells
Distance from bregma (mm)
#
i
Number of c-fos+/slc17a7+ cells
**
j
Number of c-fos+/gad2+ cells
*
k
Number of slc17a7 positive cells
l
Number of gad2 positive cells
m
Slc17a7 Gad2 c-fos

**Figure S3**. **a-b**. Single activation of the BLA-ACC pathway did not alter spontaneous locomotor activity (a; ctrl n=8; 1480±53.37; stim n=7; 1329±85.36; p=0.15), or induce avoidance in the real time avoidance test (b; RTA, ctrl: n=8; stim: n=8; $F_{(1, 14)}$=0.006; p=0.94). **c-d**. Single activation of the BLA-ACC pathway did not change anxiety-like behaviors in the NSF (c; ctrl n=8; 92.25±7.57; stim n=7; 89.14±10.97; p=0.82), or the grooming time in the ST (d; ctrl n=8; 95.25±15.13; stim n=7; 72.14±11.85; p=0.26). **e**. Repeated activation of the BLA-ACC pathway did not change spontaneous locomotor activity (3 stim: ctrl: n=8; 2281±78.89; stim: n=10; 2202±88.73; p=0.53; 6 stim: ctrl: n=8; 1812±80.24; stim: n=8; 1849±129.3; p=0.81; 9 stim: ctrl: n=8; 1656±128.0; stim: n=5; 2011±393.4; p=0.33). **f-g**. One week after the ninth stimulation no further deficit in grooming (f; ctrl: n=7; 66.57±4.86; stim: n=11; 56.55±6.22; p=0.27) or nesting behaviors (g; ctrl: n=7; stim: n=11; Chi-square=1.469; p=0.83) was observed in stimulated animals. **h**. Optogenetic activation of the BLA-ACC pathway increased c-Fos immunoreactivity in whole ACC (24a/24b) (ctrl: n=4; stim: n=4; $F_{(1, 6)}$ =17.24; p=0.006). **i-j** Optogenetic activation of the BLA-ACC pathway increased the number of *c-fos*+/*Slc17a7*+ cells (ctrl: n=5; 131.0±12.81; stim: n=5; 259.9±33.83 p=0.007) and the number of *c-fos*+/*Gad2*+ cells in the ACC (ctrl: n=5; 2.96±1.20 stim: n=5; 10.92±3.38 p=0.032). **k-l**. The number of *Slc17a7*+ (ctrl: n=5; 1136±167.0 stim: n=5; 1202±119.1, p=0.84) and *Gad2*+ cells (ctrl: n=5; 123.5±23.15 stim: n=5; 134.9±25.49 p>0.99) remained unaffected by optogenetic stimulation of the BLA-ACC pathway. **m.** Close-up image showing co-localization between *c-fos* and *Slca17a7* (green arrow) and between *c-fos* and *Gad2* (red arrow). Scale bar=20µm. Data are represented as mean±SEM. *p<0.05; **p<0.01 unpaired t-test (Spontaneous activity, NSF, ST); chi-square test for trend (Nest test); Two-Way ANOVA repeated measures (RTA: Time point x Stimulation; c-Fos immunohistochemistry: Stimulation x Anteroposteriority); Mann-Whitney test (*Slc17a7, Gad2*, *c-fos*/*Slc17a7* and *c-fos*/*Gad2* mRNA quantification).

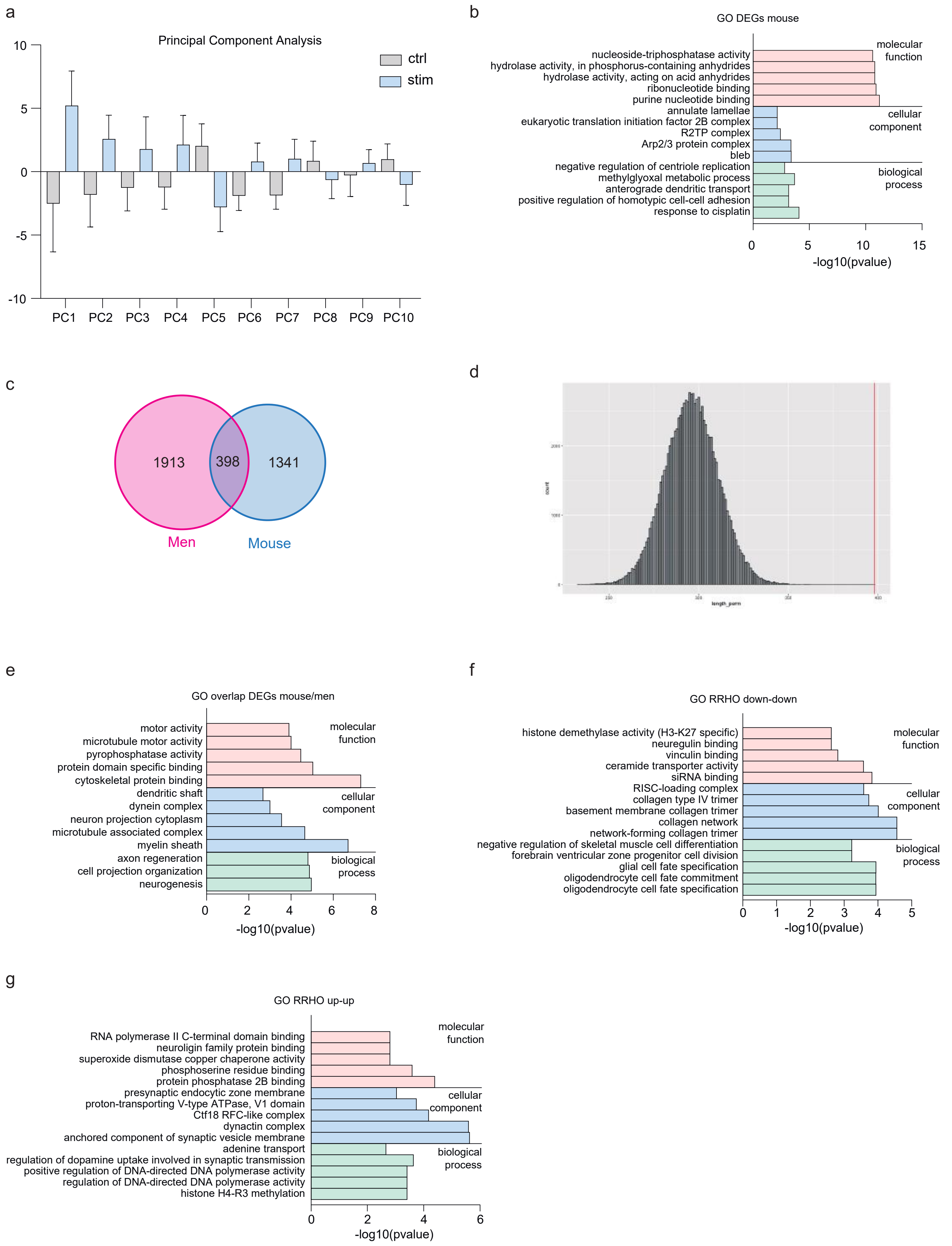

a
Principal Component Analysis
ctrl
stim
10
5
0
-5
-10
PC1 PC2 PC3 PC4 PC5 PC6 PC7 PC8 PC9 PC10
b
GO DEGs mouse
nucleoside-triphosphatase activity
hydrolase activity, in phosphorus-containing anhydrides
hydrolase activity, acting on acid anhydrides
ribonucleotide binding
purine nucleotide binding
annulate lamellae
eukaryotic translation initiation factor 2B complex
R2TP complex
Arp2/3 protein complex
bleb
negative regulation of centriole replication
methylglyoxal metabolic process
anterograde dendritic transport
positive regulation of homotypic cell-cell adhesion
response to cisplatin
molecular function
cellular component
biological process
0 5 10 15
-log10(pvalue)
c
1913
398
1341
Men
Mouse
d
length_perm
e
GO overlap DEGs mouse/men
motor activity
microtubule motor activity
pyrophosphatase activity
protein domain specific binding
cytoskeletal protein binding
dendritic shaft
dynein complex
neuron projection cytoplasm
microtubule associated complex
myelin sheath
axon regeneration
cell projection organization
neurogenesis
molecular function
cellular component
biological process
0 2 4 6 8
-log10(pvalue)
f
GO RRHO down-down
histone demethylase activity (H3-K27 specific)
neuregulin binding
vinculin binding
ceramide transporter activity
siRNA binding
RISC-loading complex
collagen type IV trimer
basement membrane collagen trimer
collagen network
network-forming collagen trimer
negative regulation of skeletal muscle cell differentiation
forebrain ventricular zone progenitor cell division
glial cell fate specification
oligodendrocyte cell fate commitment
oligodendrocyte cell fate specification
molecular function
cellular component
biological process
0 1 2 3 4 5
-log10(pvalue)
g
GO RRHO up-up
RNA polymerase II C-terminal domain binding
neuroligin family protein binding
superoxide dismutase copper chaperone activity
phosphoserine residue binding
protein phosphatase 2B binding
presynaptic endocytic zone membrane
proton-transporting V-type ATPase, V1 domain
Ctf18 RFC-like complex
dynactin complex
anchored component of synaptic vesicle membrane
adenine transport
regulation of dopamine uptake involved in synaptic transmission
positive regulation of DNA-directed DNA polymerase activity
regulation of DNA-directed DNA polymerase activity
histone H4-R3 methylation
molecular function
cellular component
biological process
0 2 4 6
-log10(pvalue)

**Figure S4**. **a**. Principal Components Analysis showed robust differences between control and stimulated animals at the whole genome level. **b**. GO enrichment (biological process, cellular component and molecular function) of the 2611 significantly changed genes in stimulated animals. **c**. Among the 1341 orthologous genes differentially expressed in mice, 398 (29.6%) were also differentially expressed in men. **d**. Expected frequency of each possible size for the interaction between mouse and men DEGs. The intersection size obtained here (398 genes) is represented by the red vertical line. **e**. GO enrichment (biological process, cellular component and molecular function) of the 398 differentially expressed genes in both mice and men. **f-g**. GO enrichment (biological process, cellular component and molecular function) of the genes commonly down-regulated (f) and up-regulated (g) in mice and men.

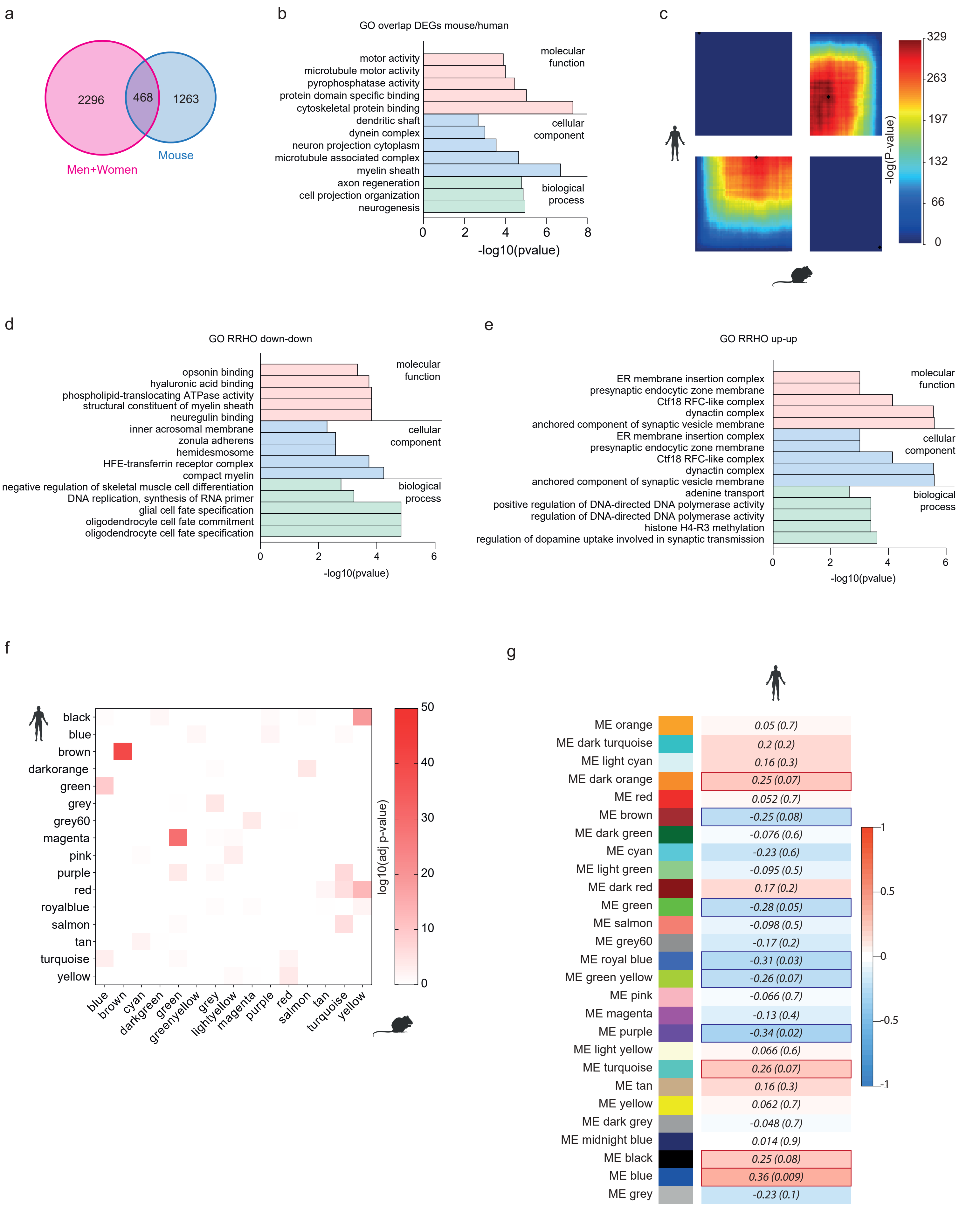

a
2296
468
1263
Men+Women
Mouse
b
GO overlap DEGs mouse/human
motor activity
microtubule motor activity
pyrophosphatase activity
protein domain specific binding
cytoskeletal protein binding
dendritic shaft
dynein complex
neuron projection cytoplasm
microtubule associated complex
myelin sheath
axon regeneration
cell projection organization
neurogenesis
molecular function
cellular component
biological process
0 2 4 6 8
-log10(pvalue)
c
329
263
197
132
66
0
-log(P-value)
d
GO RRHO down-down
opsonin binding
hyaluronic acid binding
phospholipid-translocating ATPase activity
structural constituent of myelin sheath
neuregulin binding
inner acrosomal membrane
zonula adherens
hemidesmosome
HFE-transferrin receptor complex
compact myelin
negative regulation of skeletal muscle cell differentiation
DNA replication, synthesis of RNA primer
glial cell fate specification
oligodendrocyte cell fate commitment
oligodendrocyte cell fate specification
molecular function
cellular component
biological process
0 2 4 6
-log10(pvalue)
e
GO RRHO up-up
ER membrane insertion complex
presynaptic endocytic zone membrane
Ctf18 RFC-like complex
dynactin complex
anchored component of synaptic vesicle membrane
ER membrane insertion complex
presynaptic endocytic zone membrane
Ctf18 RFC-like complex
dynactin complex
anchored component of synaptic vesicle membrane
adenine transport
positive regulation of DNA-directed DNA polymerase activity
regulation of DNA-directed DNA polymerase activity
histone H4-R3 methylation
regulation of dopamine uptake involved in synaptic transmission
molecular function
cellular component
biological process
0 2 4 6
-log10(pvalue)
f
black
blue
brown
darkorange
green
grey
grey60
magenta
pink
purple
red
royalblue
salmon
tan
turquoise
yellow
blue
brown
cyan
darkgreen
green
greenyellow
grey
lightyellow
magenta
purple
red
salmon
tan
turquoise
yellow
log10(adj p-value)
50
40
30
20
10
0
g
ME orange 0.05 (0.7)
ME dark turquoise 0.2 (0.2)
ME light cyan 0.16 (0.3)
ME dark orange 0.25 (0.07)
ME red 0.052 (0.7)
ME brown -0.25 (0.08)
ME dark green -0.076 (0.6)
ME cyan -0.23 (0.6)
ME light green -0.095 (0.5)
ME dark red 0.17 (0.2)
ME green -0.28 (0.05)
ME salmon -0.098 (0.5)
ME grey60 -0.17 (0.2)
ME royal blue -0.31 (0.03)
ME green yellow -0.26 (0.07)
ME pink -0.066 (0.7)
ME magenta -0.13 (0.4)
ME purple -0.34 (0.02)
ME light yellow 0.066 (0.6)
ME turquoise 0.26 (0.07)
ME tan 0.16 (0.3)
ME yellow 0.062 (0.7)
ME dark grey -0.048 (0.7)
ME midnight blue 0.014 (0.9)
ME black 0.25 (0.08)
ME blue 0.36 (0.009)
ME grey -0.23 (0.1)
1
0.5
0
-0.5
-1

**Figure S5. a**. Among the 1341 orthologous genes differentially expressed in mice, 468 (34.9%) were also differentially expressed in human (men+women). **b**. GO enrichment (biological process, cellular component and molecular function) of the 468 differentially expressed genes in both mice and men. **c**. Rank-Rank Hypergeometric Overlap (RRHO2) unraveled shared transcriptomic changes in the ACC across mice and human (men and women included) as a function of optogenetic stimulation (mouse) or a diagnosis of major depressive disorder (MDD). Levels of significance for the rank overlap between human and mice are color-coded, with a maximal Fisher's Exact Test p=4.69E-140 for up-regulated gene (bottom-left panel) and maximal FET p=1.62E-130 for down-regulated genes (upper-right panel). **d-e**. GO enrichment (biological process, cellular component and molecular function) of the genes commonly down-regulated (f) and up-regulated (g) in mice and human. **f**. Heatmap representing the level of significance of overlaps between mice and men gene modules (measured using the FET). The highest overlap (p=3.52E-52) was obtained for the human/brown and mouse/brown modules. **g**. WGCNA was used to analyze network and modular gene co-expression in the human ACC. The tables depict associations between individual gene modules and MDD diagnosis in human. Each row corresponds to correlations and p-values obtained against each module's eigengene.

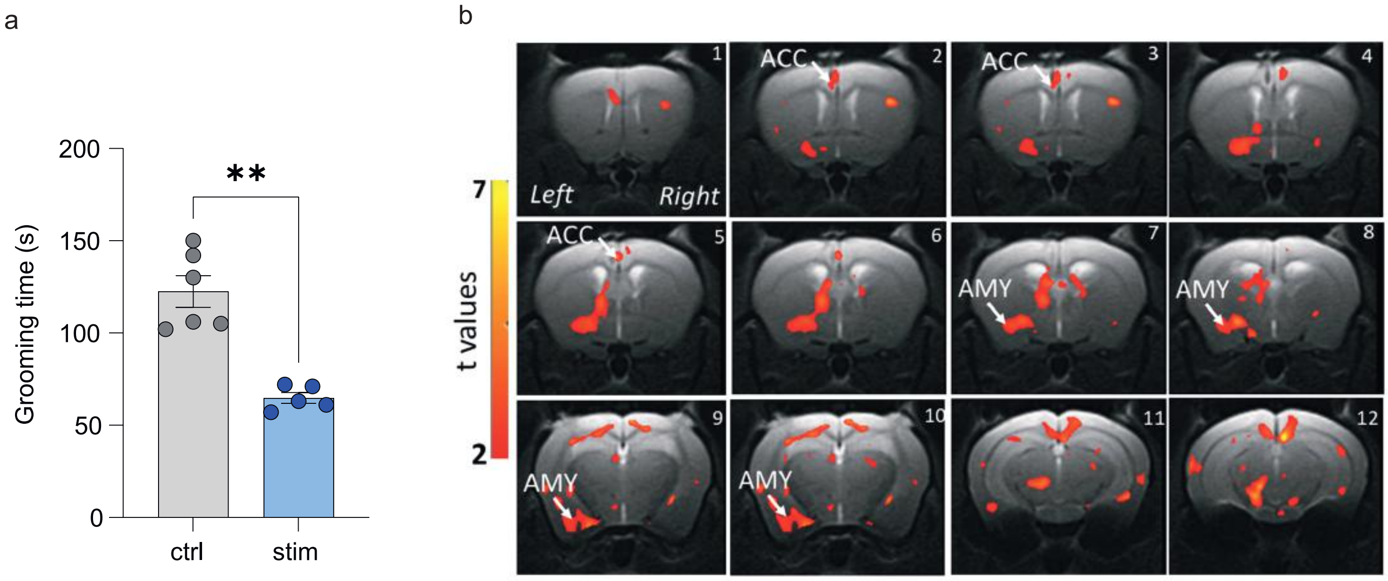
a
Grooming time (s)
200
150
100
50
0
**
ctrl
stim
b
7
t values
2
Left
Right
ACC
AMY
1
2
3
4
5
6
7
8
9
10
11
12

**Figure S6. a**. Repeated activation of the BLA-ACC decreases grooming time in stimulated animals (ctrl n=6; 122.5±8.55; stim n=5; 64.80±2.90; p<0.002). **b**. Representative coronal MRI images showing in red the brain areas with significant positive correlation between fractional anisotropy values and grooming time (ctrl n=8; stim=7; p<0.001; uncorrected). Microstructural changes reflected by low fractional anisotropy (FA) along the left BLA-ACC pathway (see Fig. 6d) correlate with low grooming time in stimulated animals. Data are represented as mean±SEM. **p<0.01. unpaired t-test Mann-Whitney test (ST).

**Supplementary Table 2: Human and Mouse cohort metrics**

| **Human** | **MDD** (n=26) | **Controls** (n=24) |
|---|---|---|
| Age (Years) | 42,2 ± 2.85 | 46.2 ± 4.4 |
| Sex (M/F) | 19/7 | 19/5 |
| RIN | 6.83 ± 0.15 | 6.86 ± 0.12 |
| **Mouse** | **Stimulated** (n=10) | **Controls** (n=12) |
| Batch (1/2) | 4/6 | 6/6 |
| RIN | 9.40 ± 0.28 | 9.44 ± 0.15 |

**Supplementary Table 7 : GO term enriched in the 5 mouse modules significantly associated with optogenetic stimulation and conserved in human**

| module | enrichmentP | BonferoniP | termOntology | termName |
|---|---|---|---|---|
| **blue** | 1,49E-08 | 0,000269653 | CC | preribosome |
| **blue** | 1,55E-06 | 0,028138839 | BP | RNA processing |
| **blue** | 3,32E-06 | 0,060160709 | CC | cytoplasmic vesicle |
| **blue** | 3,82E-06 | 0,069182237 | CC | intracellular vesicle |
| **blue** | 8,64E-06 | 0,156507228 | CC | clathrin-coated vesicle membrane |
| **blue** | 9,72E-06 | 0,175981124 | CC | clathrin-coated vesicle |
| **blue** | 9,94E-06 | 0,179982376 | CC | protein-containing complex |
| **blue** | 1,08E-05 | 0,195746883 | BP | mRNA processing |
| **blue** | 1,19E-05 | 0,215232033 | MF | RNA binding |
| **blue** | 1,25E-05 | 0,226475177 | MF | ATP-dependent protein binding |
| **brown** | 2,89E-10 | 5,23E-06 | BP | myelination |
| **brown** | 4,61E-10 | 8,36E-06 | BP | axon ensheathment |
| **brown** | 1,39E-08 | 0,000252159 | CC | myelin sheath |
| **brown** | 7,71E-08 | 0,001395487 | BP | oligodendrocyte differentiation |
| **brown** | 8,93E-08 | 0,001616898 | BP | glial cell differentiation |
| **brown** | 9,37E-08 | 0,001697005 | BP | gliogenesis |
| **brown** | 1,80E-07 | 0,003260951 | BP | nervous system development |
| **brown** | 1,87E-07 | 0,003391081 | CC | cell periphery |
| **brown** | 3,49E-07 | 0,006325485 | BP | neurogenesis |
| **brown** | 1,21E-06 | 0,02197177 | CC | plasma membrane |
| **greenyellow** | 0,00015471 | 1 | BP | neuronal action potential |
| **greenyellow** | 0,00036253 | 1 | MF | ion channel activity |
| **greenyellow** | 0,00059938 | 1 | MF | channel activity |
| **greenyellow** | 0,000614 | 1 | MF | passive transmembrane transporter activity |
| **greenyellow** | 0,00069401 | 1 | BP | action potential |
| **greenyellow** | 0,00084187 | 1 | MF | ion transmembrane transporter activity |
| **greenyellow** | 0,00094943 | 1 | BP | trigeminal nerve structural organization |
| **greenyellow** | 0,00103757 | 1 | MF | cation channel activity |
| **greenyellow** | 0,00126358 | 1 | CC | calyx of Held |
| **greenyellow** | 0,00141493 | 1 | BP | trigeminal nerve development |

| | | | | |
|---|---|---|---|---|
| **magenta** | 0,0002393 | 1 | MF | ubiquitin-like protein-specific protease activity |
| **magenta** | 0,00082264 | 1 | CC | nucleoplasm |
| **magenta** | 0,00118341 | 1 | MF | nucleic acid binding |
| **magenta** | 0,00218795 | 1 | BP | protein desumoylation |
| **magenta** | 0,00228823 | 1 | MF | cysteine-type peptidase activity |
| **magenta** | 0,00255046 | 1 | CC | cytoplasmic stress granule |
| **magenta** | 0,00349927 | 1 | CC | nucleus |
| **magenta** | 0,0038129 | 1 | CC | nuclear lumen |
| **magenta** | 0,0045933 | 1 | BP | double-strand break repair via break-induced replication |
| **magenta** | 0,00480606 | 1 | BP | lung alveolus development |
| **yellow** | 1,90E-15 | 3,44E-11 | BP | mitochondrial translational termination |
| **yellow** | 5,13E-15 | 9,28E-11 | BP | translational elongation |
| **yellow** | 5,24E-15 | 9,49E-11 | CC | mitochondrial inner membrane |
| **yellow** | 1,05E-14 | 1,90E-10 | BP | mitochondrial translational elongation |
| **yellow** | 1,51E-13 | 2,74E-09 | BP | translational termination |
| **yellow** | 2,07E-13 | 3,74E-09 | BP | mitochondrial translation |
| **yellow** | 2,58E-13 | 4,67E-09 | CC | mitochondrial envelope |
| **yellow** | 3,71E-13 | 6,72E-09 | CC | organelle inner membrane |
| **yellow** | 9,12E-13 | 1,65E-08 | CC | mitochondrial membrane |
| **yellow** | 1,54E-12 | 2,79E-08 | BP | regulation of cellular amino acid metabolic process |

**Supplementary Table 8: Primers sequences for RT-qPCR**

| Gene name | Sequence |
|---|---|
| *B2m* | F TGCTACGTAACACAGTTCCACC<br>R: TCTGCAGGCGTATGTATCAGTC |
| *Gapdh* | F: TGGCCTCCAAGGAGTAAGAAAC<br>R: TGGGATGGAAATTGTGAGGGAG |
| *Actb* | F: ATCAGCAAGCAGGAGTACGATG<br>R: GGTGTAAAACGCAGCTCAGTAAC |
| *Gusb* | F: TACCGACATGAGAGTGGTGTTG<br>R: TAATGTCAGCCTCAAAGGGGAG |
| *Plp1* | F: AGCAAAGTCAGCCGCAAAAC<br>R: TGAGAGCTTCATGTCCACATCC |
| *Ermn* | F: AGAGAACCTCTTCGTTGTTCACC<br>R: TTGCTGGGCAGTTCTTTCCTTC |
| *Aspa* | F: CACTTCTAACATGGGTTGCACTC<br>R: AAACAGAGCAGGGTAATGGAGC |
| *Ugt8* | F: TGAAGGAGAGCTGTATGATGCC<br>R: TCCGTCATGGCGAAGAATGTAG |
| *Mal* | F: ATTACCATGAAAACATCGCCGC<br>R: TTAATGGGGAAGATGGGCTGAC |
| *Mog* | F: CATAAAGATGGCCTGTTTGTGGAG<br>R: CCCTGGTCCTATCACTCTGAATTG |
| *Mbp* | F: ATCGGCTCACAAGGGATTCAAG<br>R: TATATTAAGAAGCCGAGGGCAGG |
| *Lingo1* | F: CAACAAGACCTTCGCCTTCATC<br>R: TGCTTTGTGTTGCCTTTGCC |
| *Sema4a* | F: AGCCATGTGGTCATGTATCTGG<br>R: TGAATCTCCTCCACGAGATAAGC |
